# PRISM: Niche-informed Deciphering of Incomplete Spatial Multi-Omics Data

**DOI:** 10.64898/2026.02.03.703456

**Authors:** Shiguan Mu, Zhikang Wang, Yi Liao, Jiaming Liang, Daoliang Zhang, Chuyao Wang, Jiahui Xie, Xiaoqi Sheng, Tinghe Zhang, Weitian Huang, Jiangning Song, Zhiyuan Yuan, Hongmin Cai

## Abstract

Spatial multi-omics data, characterizing the knowledge of diverse molecular layers, have become indispensable for the *in situ* analysis of tissue architecture and complex biological processes. Nevertheless, current spatial multi-omics sequencing protocols are often hindered by incompatible protocols, resulting in incomplete spatial multi-omics pairing due to inconsistent field-of-view or spatial resolution. To address this, we present PRISM, a novel computational method tailored for this scenario. PRISM leverages a niche-informed graph to propagate information from paired to unpaired regions, jointly achieving the spatial domain identification and spatial omics imputation. Extensive benchmarking on five diverse simulated and real experimental datasets demonstrated that PRISM outperformed existing methods in spatial multi-omics analysis tasks. Application to the human Parkinson’s disease data revealed that PRISM accurately recovered dopamine-associated spatial domains and metabolite distributions masked by incomplete data gaps. PRISM offers a robust solution for bridging the integration gap inherent in incompatible sequencing protocols, thereby facilitating more accurate downstream biological interpretation.

## 1. INTRODUCTION

Spatial multi-omics technologies provide an *in situ* perspective of tissues by capturing multiple molecular layers while preserving native tissue architecture^1-3^. This multi-modal phenotypic characterization of diverse molecular layers reveals the fundamental principles governing tissue organization within its spatial context^4^. For instance, spatial CITE-seq co-profiles transcripts and proteins to uncover compartment-specific immune programs elusive to single-modality analysis^5,6^. Similarly, spatial multi-modal analysis (SMA) protocol integrates transcriptomics with metabolomics to reveal spatially organized metabolic patterns and alterations relevant to Parkinson’s disease (PD)^7^. Furthermore, co-profiling gene expression with chromatin landscapes via MISAR-seq^8^ or spatial CUT&Tag-RNA-seq^9^ connects epigenetic regulatory dynamics to developmental cell-fate decisions. By synthesizing these complementary insights, cross-platform multi-modal atlases are increasingly refining functional tissue zonation and compartmentalization at scale^10-12^.

Spatial multi-omics datasets are typically generated under two major acquisition paradigms^13-19^. The first paradigm utilizes co-profiling strategies that ensure intrinsic spatial correspondence^8,9^. By employing a unified spatial indexing system, these methods simultaneously barcode multiple modalities within the same physical pixel. Representative methodologies include MISAR-seq^8^ for simultaneous chromatin accessibility (via assay for transposase-accessible chromatin, or ATAC) and gene expression, alongside Stereo-CITE-seq^15^ for joint RNA and protein detection. This paradigm has recently advanced to spatial tri-omics, ensuring precise bin-level alignment across chromatin, transcriptomic, and proteomic layers^20^. Limited accessibility and insufficient commercialization frequently restrict the application of these advanced co-profiling techniques to specialized research laboratories. In contrast, an alternative paradigm integrates multiple incompatible assays either from a single tissue section or adjacent sections into a unified coordinate system^17,21-24^. These datasets frequently exhibit inconsistent fields of view (FOV) or spatial resolution^24,25^. Lacking an inherent shared coordinate system, these sequential measurements necessitate post-hoc spatial registration to align spatial correspondence^26-30^. This practice is common when combining widely utilized commercial workflows such as the integration of 10x Visium transcriptomics with CODEX imaging for high-plex protein detection or with MALDI mass spectrometry imaging for localized metabolic profiling^31,32^.

Several computational methods have been developed to integrate and analyze fully registered tissue sections^33,34^. SpatialGlue^11^ applies dual-graph contrastive learning to identify spatial domains by learning joint representations from co-located modalities, while COSMOS^35^ and MISO^36^ employ cross-modal alignment and whole-slide image integration, respectively, to facilitate multimodal analysis. Additionally, SpaMosaic^37^ enables the stitching of multiple tissue sections through latent-space anchoring. However, all existing approaches were developed based on fully registered spatial multi-omics data. This often fails to account for incompletely registered data resulting from technical limitations inherent in the acquisition process^38-40^.

Here we present PRISM, an integrative computational framework designed for spatial multi-omics under such incomplete registration. Unlike existing methods requiring fully registered inputs, PRISM seamlessly interfaces with alignment workflows (e.g., MAGPIE^41^ and SLAT^42^) to facilitate robust spatial domain identification and spatial omics imputation on incomplete spatial multi-omics data. Built on a spatial niche similarity hypothesis^43^, PRISM assumes that cells or spatial spots within comparable local spatial contexts share congruent cross-modal profiles. To operationalize this principle, PRISM utilizes niche-informed graphs within a unified architecture of transformers^44^ and graph attention networks^45^. This framework propagates information from registered to unregistered cells to learn robust multi-modal representations, effectively equipping every individual cell with a comprehensive multi-omics profile. We assessed PRISM on representative tissues, platforms and modality pairs that recapitulate real-world acquisition scenarios^7-9,21^. These included FOV-induced incompleteness in human lymphoid organs (RNA-protein) and embryonic mouse brains (RNA-ATAC), as well as cross-platform integration exhibiting resolution-induced incompleteness in human PD striatum (RNA-metabolomics). Furthermore, we applied PRISM to a hybrid incompleteness scenario involving adjacent postnatal day 22 (P22) mouse brain sections, where histone modifications (H3k27ac) and RNA profiles were subject to simultaneous inconsistent FOV and spatial resolution. Notably, PRISM restored the spatial continuity of dopaminergic functional domains and metabolic patterns, uncovering biological insights obscured by incomplete spatial multi-omics data. Across these diverse datasets, PRISM improved spatial domain identification, spatial omics imputation, and tissue-level continuity relative to existing spatial multi-omics^11,35-37^ and single-cell translation methods^46-48^.

## 2. RESULTS

### Motivation

Spatial multi-omics technologies provide a transformative perspective on tissue architecture by mapping multi-layered molecular information while preserving native spatial context^49-51^. Based on prevalent acquisition strategies (Figure 1a), we categorized protocols into two distinct regimes: compatible and incompatible protocols (Figure 1b, c). This strategy is defined by whether disparate molecular modalities are captured within an intrinsic spatial coordinate system or necessitate post-hoc registration to align within a unified reference^52,53^.

**Fig. 1:**
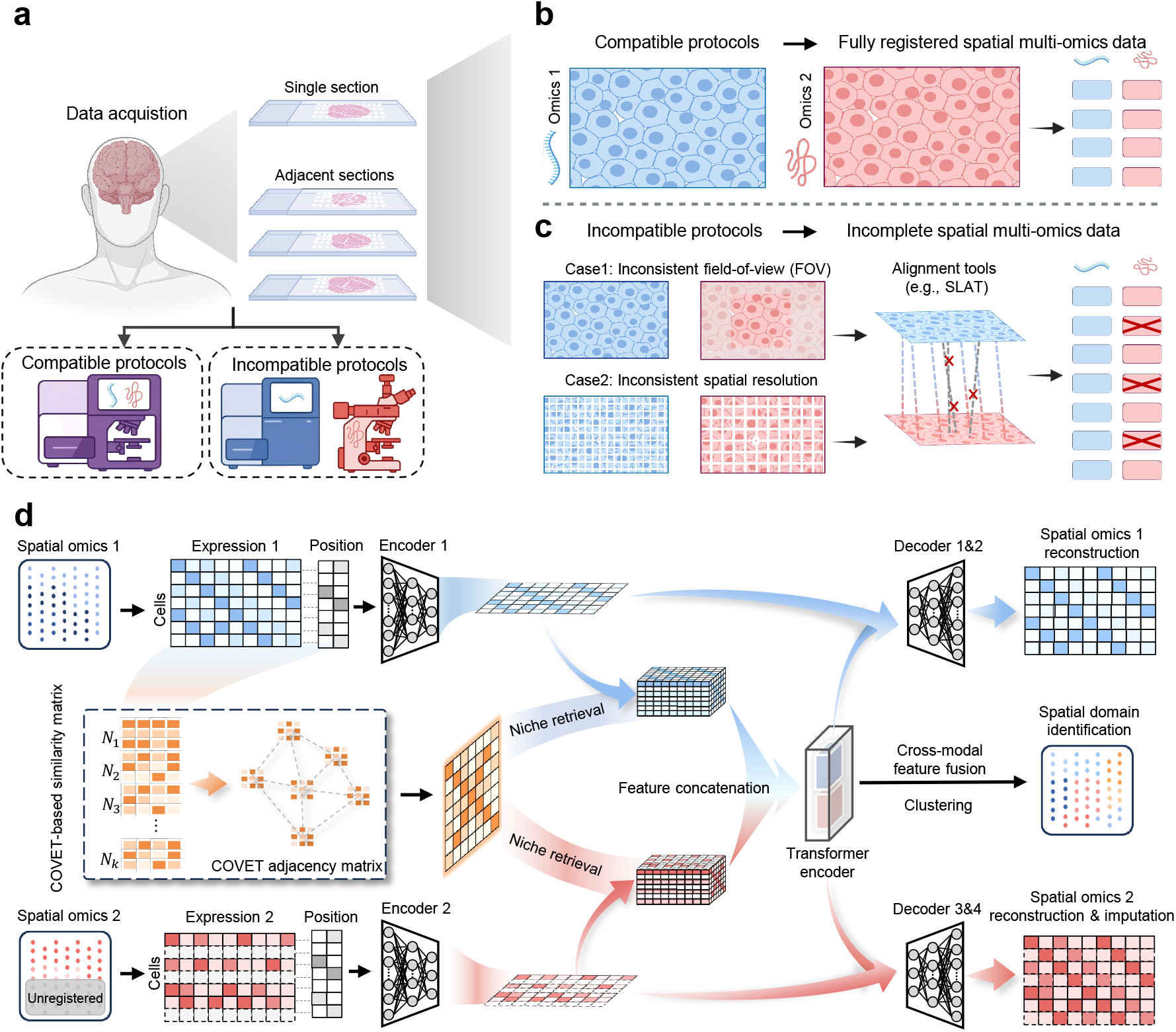
Spatial multi-omics data acquisition under compatible and incompatible protocols and PRISM overview. **a**, Spatial multi-omics acquisition from the single section (co-assay) or adjacent sections, often using compatible or incompatible protocols. **b**, Ideal case with compatible protocols producing fully registered multi-omics observations. **c**, Practical cases utilizing incompatible protocols lead to incomplete registeration, characterized by inconsistent FOV or spatial resolution. Alignment tools (e.g., SLAT) then yield incomplete spatial multi-omics data after filtering low-confidence matches. **d**, PRISM architecture: modality-specific encoders, COVET-based niche graph for retrieval and propagation, transformer-based cross-modal fusion, and decoders for reconstruction/imputation with downstream spatial domain identification.

Compatible protocols achieve intrinsic spatial correspondence by co-profiling multiple molecular layers within a unified spatial indexing system^20^. Beyond limited accessibility, these assays frequently encounter inherent biochemical conflicts where optimizing sensitivity for one modality often compromises the resolution of others. Specifically, the intense permeabilization required for epigenomic nuclear access inevitably triggers cytoplasmic mRNA leakage, degrading transcriptomic resolution ^9^. Similarly, fixatives optimized for protein preservation often inhibit enzymatic activities essential for nucleic acid detection ^3,54^. Ultimately, these antagonistic requirements impose a performance ceiling, forcing a persistent compromise between modality-specific sensitivity and spatial fidelity.

In contrast, incompatible protocols represent a more widely adopted paradigm due to the commercial maturity of platforms such as 10x Visium and Xenium^17^. This approach encompasses both single-section multi-omics and adjacent-sections single-omics^53,55^, both of which face substantial technical hurdles. In single-section multi-omics, sequential assays often encounter disparate spatial grids characterized by inconsistent FOV or spatial resolution^7^. These discrepancies arise from hardware limits or the aggregation of sparse signals to ensure stability, leading to misaligned molecular layers and mismatched spatial coverage^41^. Conversely, adjacent-section single-omics is inherently more complex because physical tissue handling introduces non-linear morphological distortions^27^. Beyond these structural misalignments, this approach could also lead to reconcile simultaneous inconsistent FOV and spatial resolution. Collectively, these factors aggregate into a fragmented molecular landscape that complicates joint analysis and obscures underlying biological signals.

Both scenarios necessitate spatial registration to align multi-modal features within a unified coordinate system prior to joint analysis. Registration methods range from global rigid transformations (e.g., translation, rotation, scaling), which are often adequate for co-assays on the same section^52^, to non-rigid warping that is essential for adjacent sections where sectioning and processing introduce nonlinear distortions^30^. Even with careful pipelines, registration typically yields only partial spatial overlap, producing datasets that mix co-registered multi-omics locations with single-modality regions^56-58^. This incomplete structure does not fit classical vertical, horizontal or diagonal integration^37,59,60^, because only a limited subset of cross-modal features and positions can serve as anchors. Although shared latent-variable models begin to address similar issues in single-cell multi-omics^57,61-63^, dedicated computational frameworks for spatial multi-omics under such incomplete registration remain lacking.

### PRISM overview

PRISM is an adaptive spatial multi-omics integration framework built on the premise that cells or spots sharing analogous spatial contexts are likely to exhibit congruent molecular profiles. Accordingly, PRISM can simultaneously perform the spatial domain identification and spatial omics imputation on the incomplete spatial multi-omics data (Figure 1d). Specifically, PRISM first extracts similar covariance expression patterns between niches based on covariance environment (COVET) prior^43^, followed by constructs a COVET-based similarity matrix. As a key component of PRISM, it is employed to aggregate mean encoded features of similar cells/spots via niche retrieval, thereby ensuring precise spatial omics imputation within unregistered regions. Building on this structure, a transformer encoder captures cross-modal dependencies to effectively model complex interactions across distinct molecular modalities. A two-layer linear projection then derives modality-specific embeddings that preserve the intrinsic signatures of each omics layer and support faithful reconstruction. Concurrently, cross-modal feature fusion extracts their interactive representations that capture emergent spatial structures for robust spatial domain identification. Finally, two-layer graph attentional decoders leverage local spatial constraints to reconstruct measured signals and predict missing molecular profiles. This integrated workflow enables PRISM to restore spatial continuity by overcoming the systematic data missingness inherent in incompatible acquisition protocols.

We validated PRISM across two prevalent forms of incomplete spatial multi-omics data originating from incompatible acquisition protocols (Figure 1c). The first category is FOV-induced incompleteness, which occurs when distinct modalities encompass non-overlapping tissue regions. The second category is resolution-induced incompleteness, which arises when disparate sampling resolutions and grid geometries preclude direct bin-level alignment. By leveraging niche-informed priors across these benchmarks, PRISM successfully incorporated unregistered regions into a unified analytical framework. This approach consistently achieved spatial domain identification and modality reconstruction performance on par with gold-standard, fully registered multi-omics data.

### Integration of FOV-induced incomplete spatial multi-omics data from human lymphoid organs

We employed a human tonsil dataset consisting of three spatially resolved sections, with each section characterized by fully registered spatial transcriptomics and proteomics data (generated using the 10x Genomics Visium platform with Antibody-Derived Tags (ADTs)) (Figure 2a). To simulate a FOV-induced incompleteness, a contiguous 50% region of the spatial proteomics data was masked (Figure 2b, c).

**Fig. 2:**
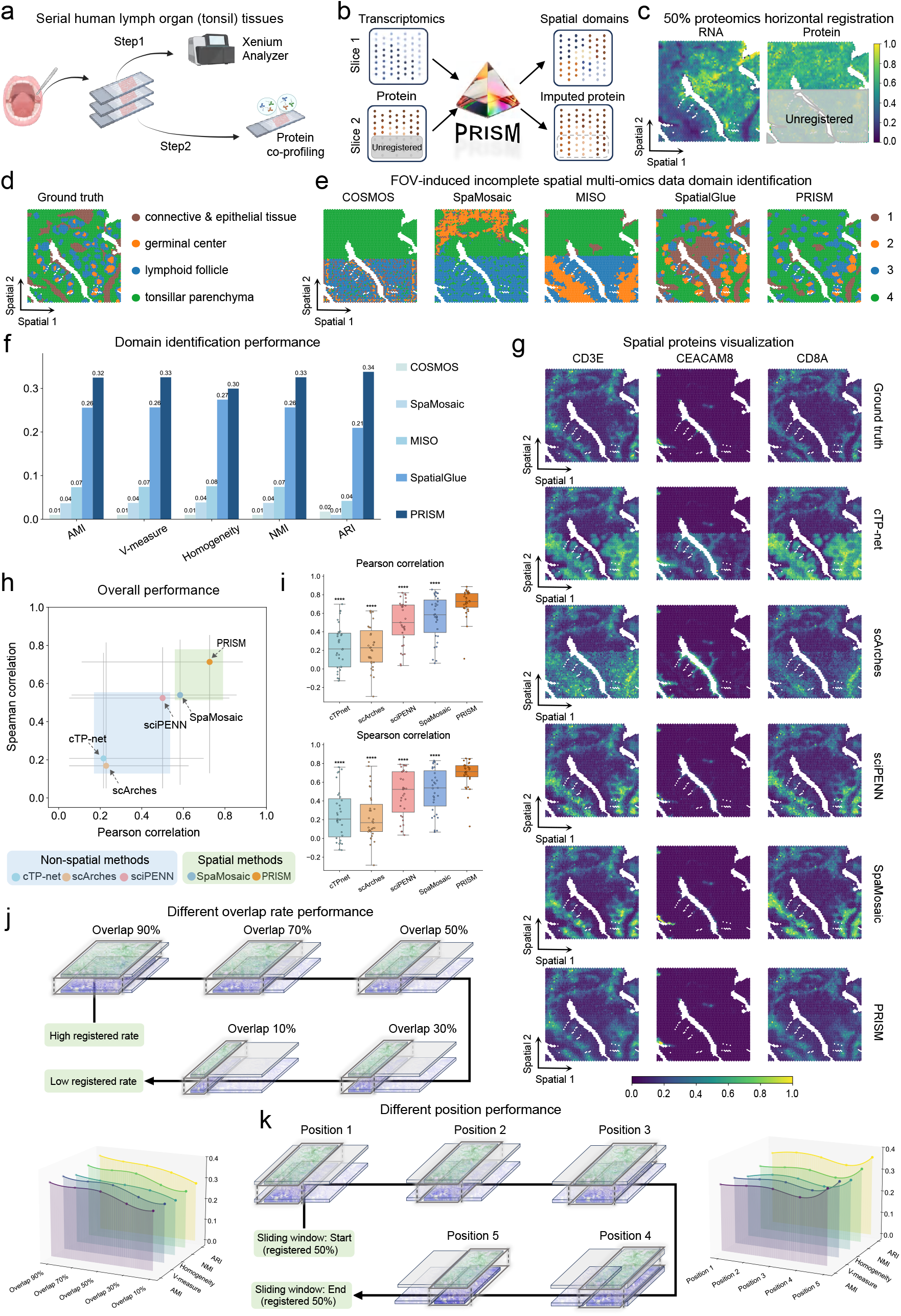
Evaluation of PRISM under FOV-induced incompleteness in human tonsil. **a**, Experimental schematic for generating spatial multi-omics data from serial human lymphoid organ (tonsil) sections, with transcriptomics profiling (Step 1) and spatial protein co-profiling (Step 2). **b**, Analysis workflow: incomplete spatial multi-omics inputs (containing an unregistered proteomics region) are processed by PRISM to identify spatial domains and impute protein signals in unregistered locations. **c**, Simulation of FOV-induced incompleteness by masking a contiguous horizontal region (50%) of the proteomics modality while retaining full transcriptomics coverage (the masked area is treated as unregistered). **d**, Ground-truth spatial domain annotation of the tonsil section (connective & epithelial tissue, germinal center, lymphoid follicle, and tonsillar parenchyma). **e**, Comparison of spatial domain identification results between PRISM and advanced spatial multi-omics baseline method. **f**, Quantitative comparison of domain identification using AMI, V-measure, homogeneity, NMI and ARI. **g**, Spatial visualization of representative proteins: ground truth measurements compared with predictions from non-spatial translators and spatial methods. **h**, Overall protein imputation performance is summarized by PCC and SPCC across methods. **i**, Compare the protein-related distributions (PCC and SPCC) of different methods using box plots. Each dot in the box represents one protein. The statistical significance of PRISM and other methods were indicated using * (*: p-value<0.05, **: p-value<0.01, ***: p-value<0.001, ****: p-value<0.0001). **j**, Robustness analysis under varying FOV overlap rates (from 90% down to 10%), illustrating performance trends as the registered region decreases. **k**, Robustness analysis regarding the spatial position of the registered window: a sliding-window scheme with fixed overlap (50%) evaluated across multiple positions.

We compared PRISM with four state-of-the-art spatial multi-omics analysis methods, i.e., SpatialGlue^11^, COSMOS^35^, MISO^36^ and SpaMosaic^37^, in terms of spatial domaining analysis. Here, the ground truth (Figure 2d) and results of the second section were particularly illustrated (Figure 2e). Notably, PRISM and SpatialGlue were the only methods to successfully recapitulate the general tissue architecture. However, only PRISM can distinguish the germinal center from the lymphoid follicle, critical niches for B-cell maturation and adaptive immunity. This is attributed to the unique design of PRISM using niche-informed signal propagation. Apart from the qualitative visualization, quantitative assessments were conducted using five established clustering metrics: Adjusted Mutual Information (AMI), V-measure, Homogeneity, Normalized Mutual Information (NMI), and Adjusted Rand Index (ARI). Quantitative assessment confirmed that PRISM consistently ranked first across all five metrics, surpassing the second-best method by 12.85% in ARI and 6.90% in NMI (Figure 2f). Identical phenotypes and conclusions can be drawn on remaining sections (Supplementary Figure 1a-c and 2a-c).

Imputation is another key capability of PRISM. Here, we utilized the Pearson Correlation Coefficient (PCC) and Spearman Correlation Coefficient (SPCC) for performance evaluation (Figure 2h). For comprehensive benchmarking, PRISM was compared not only against the state-of-the-art spatial multi-omics method SpaMosaic but also against three leading single-cell multi-omics translation techniques, cTP-net^46^, scArches^47^, and sciPENN^48^. PRISM achieved the highest mean PCC and SPCC (Figure 2h), and per-target rankings confirmed a significant advantage over all baselines (Figure 2i). These results further highlight the critical role of spatial information during the imputation process. In Figure 2g, we visualized three representative proteins, i.e., *CD8A, CEACAM8* and *CD3E*. It was apparent that the results from single-cell translation methods have a clear artifact between the translated and measured regions. In terms of the spatial methods, the results of PRISM demonstrated a higher degree of fidelity, i.e., more similar to the measurements.

We further assessed the robustness of PRISM under FOV-induced incompleteness by systematically varying two factors: (i) the overlapping rate of two FOVs and (ii) the spatial position of the registered window. First, we simulated five distinct overlap levels, namely 90%, 70%, 50%, 30%, and 10% (Figure 2j). Second, we implemented a sliding-window scheme with a fixed 50% overlap rate, advancing the window position in 10% cellular increments at each step (Figure 2k). Across all settings, multiple quantitative metrics indicated that PRISM remained stable (Supplementary Figure 3a-d), highlighting its scalability and robustness to FOV-induced incomplete spatial multi-omics.

**Fig. 3:**
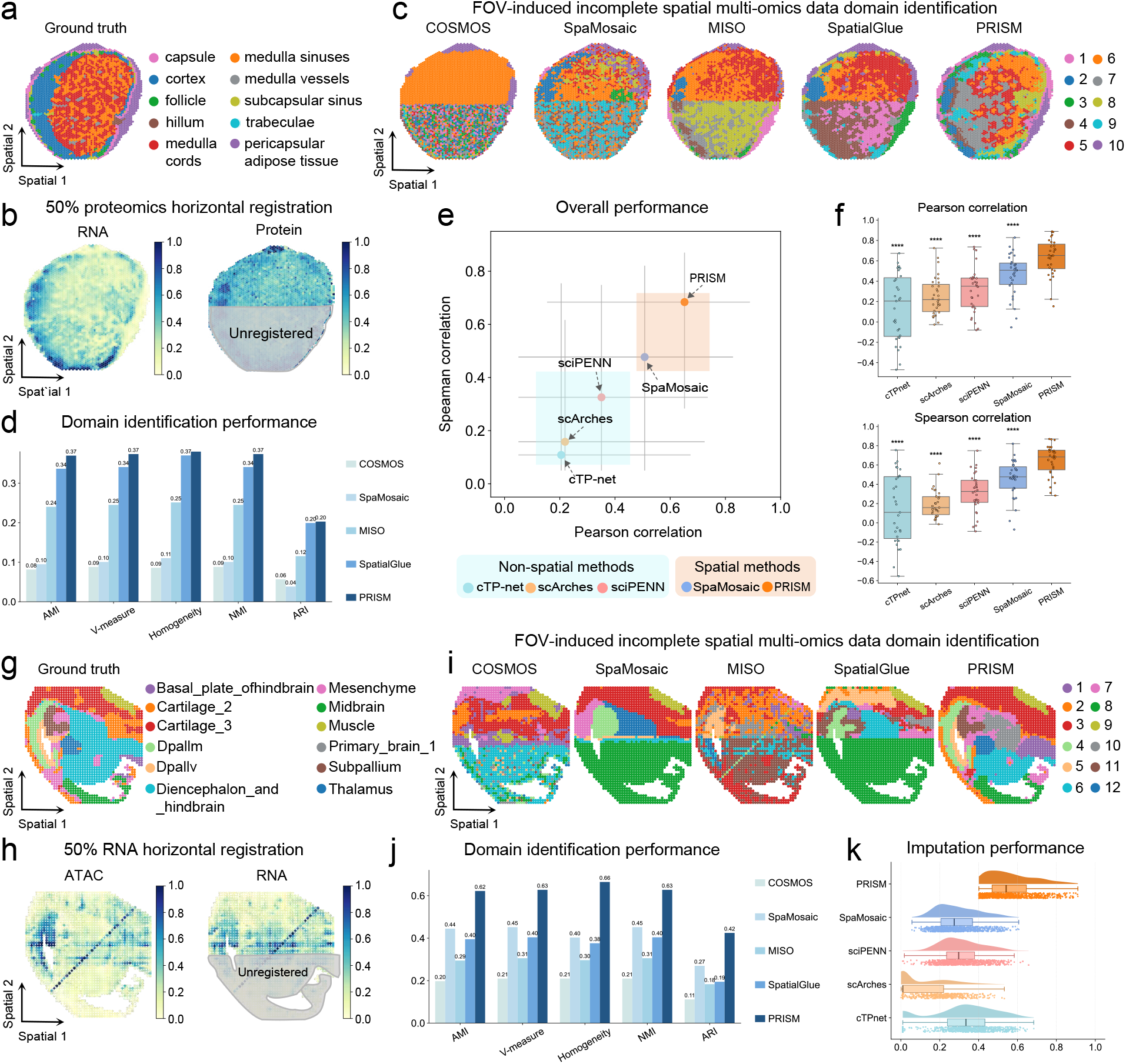
Performance of PRISM under FOV-induced incompleteness in human lymph node and mouse embryonic brain. **a**, Ground-truth spatial domain annotations for human lymph node section. **b**, Simulation of FOV-induced incompleteness in the lymph node dataset, achieved by masking a contiguous horizontal region (50%) of the protein modality while retaining full RNA coverage (simulating an incomplete spatial multi-omics scenario). **c**, Spatial domain identification under the FOV-induced incompleteness setting using state of-the-art methods. **d**, Quantitative evaluation of spatial domain identification using five established metrics. **e**, Overall imputation performance summarized by PCC and SPCC, comparing non-spatial translation baselines and spatial methods. **f**, Distribution of per-gene imputation correlations (PCC and SPCC) across methods. **g**, Ground-truth spatial domain annotations for the embryonic mouse brain section. **h**, Simulation of FOV-induced incompleteness in the embryonic brain, achieved by masking a contiguous region (50%) of the RNA modality while retaining full ATAC coverage. **i**, Visual comparison of spatial domains identified by PRISM versus baselines under the incomplete spatial multi-omics setting. **j**, Quantitative benchmarking of domain identification for the embryonic brain dataset. **k**, Imputation performance for the embryonic brain dataset shown as the distribution of correlation scores across genes, comparing PRISM with competing methods.

We next evaluated the methods on human lymph nodes, another key immune organ. The dataset consisted of three spatially resolved sections generated using the 10x Genomics Visium platform, including fully registered spatial transcriptomics and proteomics data. Ground-truth annotations delineated the canonical compartments of the lymph node (Figure 3a). Following our above setting, we masked 50% of the spatially contiguous regions in the spatial proteomics to zero while retaining the full spatial transcriptomics data (Figure 3b).

While performing the spatial domain analysis, we found that COSMOS, SpaMosaic, and MISO consistently exhibited spatial domain deviations and noise artifacts in unregistered spots (Figure 3c). While SpatialGlue achieved smoother spatial domain delineation, all four methods had discontinuities between registered and unregistered spots. In contrast, PRISM maintained superior spatial continuity across the entire tissue domain, achieving the closest registration with ground truth annotations. These results suggested that incomplete registration disrupts the model’s ability to preserve tissue continuity, a challenge PRISM overcomes through its advanced signal propagation mechanism. Quantitative evaluation (Figure 3d) revealed that PRISM outperforms all comparative methods, particularly in ARI and NMI. Identical phenotypes and conclusions can be drawn on remaining sections (Supplementary Figure 4c and 5c).

**Fig. 4:**
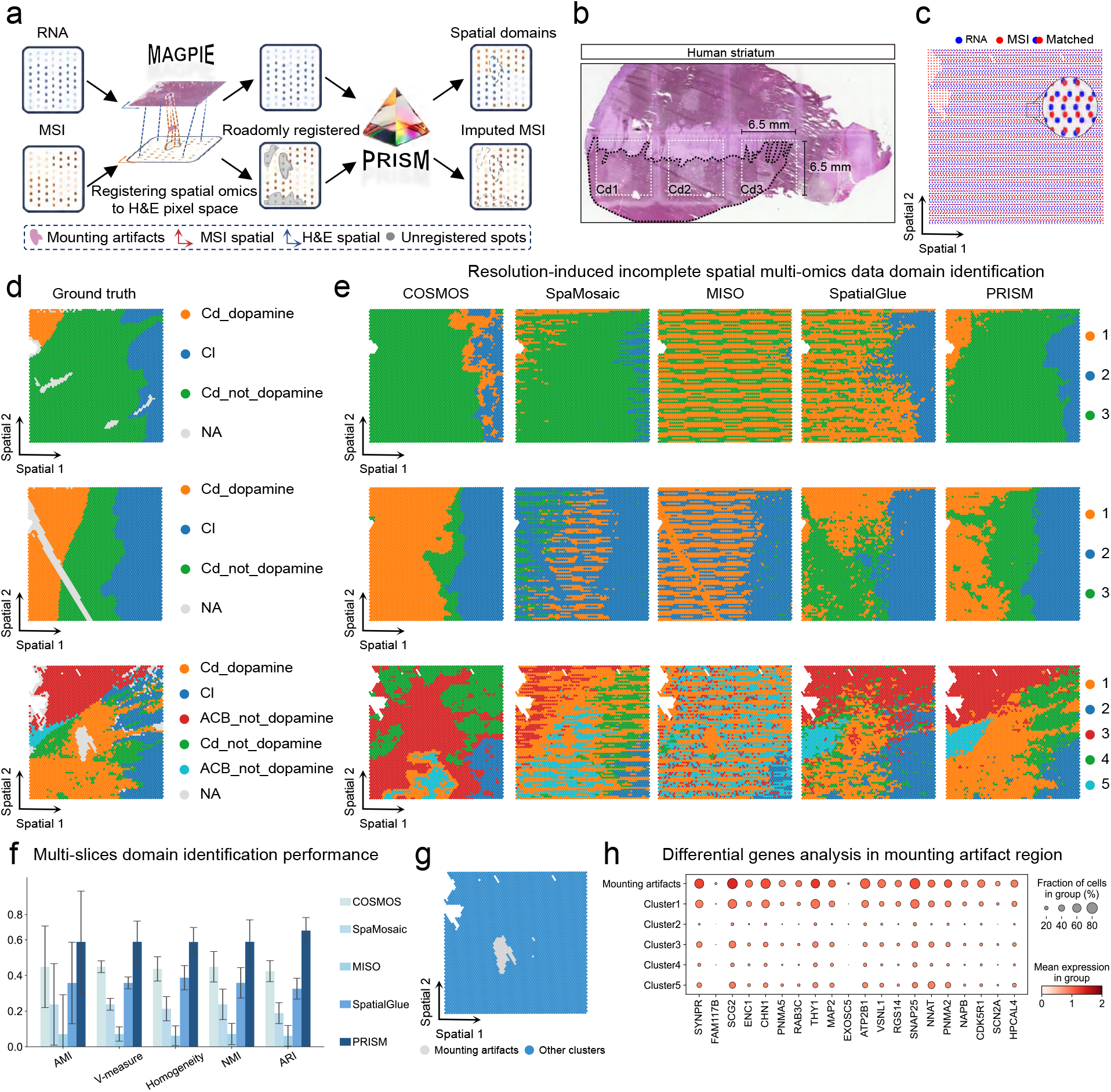
Evaluation of PRISM under resolution-induced incompleteness in human PD striatum. **a**, Workflow schematic: Visium RNA (55 µm spots) and MALDI-MSI (100 µm pixels) generated via incompatible platforms are co-registered using MAGPIE, yielding an incomplete spatial multi-omics dataset with resolution-induced gaps. PRISM then infers spatial domains and imputes MSI signals in unregistered locations. **b**, H&E image of the human striatum section with three distinct coronal regions (Cd1-Cd3) labeled. **c**, Illustration of cross-modal correspondence after registration, showing RNA spots, MSI pixels and matched pairs within the aligned coordinate system. **d**, Ground-truth spatial domain annotations for the three regions. NA: non-annotated or filtered areas (including mounting artifacts). **e**, Visual comparison of spatial domains identified by PRISM versus state-of-the-art baselines under the resolution-induced incompleteness. **f**, Quantitative benchmarking of domain identification performance across sections using five established metrics. **g**, Spatial map highlighting the mounting-artifact region and its assignment relative to other clusters. **h**, Differential gene expression summary for the mounting-artifact region versus inferred clusters, showing representative marker genes with dot size indicating the fraction of cells and color indicating mean expression.

Imputation accuracy was assessed using PCC and SPCC (Figure 3e-f). PRISM achieved the highest mean scores and showed significant improvements over each comparator (p < 0.0001) (Figure 3f). Quantitative illustration of representative proteins (*CD3E, HLA-DRA*, and *VIM*) revealed that non-spatial translators exhibited noticeable cross-boundary discontinuities typical of incomplete registration, while SpaMosaic reduced this seam but still over-expressed signals in unregistered spots. In contrast, PRISM accurately recovered ground-truth intensity ranges and intra-tissue gradients (Supplementary Figure 6a).

Under the FOV-induced incompleteness setting, PRISM significantly outperformed baselines on human lymphoid organ datasets. It achieved top-ranked spatial domain identification by quantitative metrics and was the only method to resolve critical immune niches (e.g., germinal centers), while also delivering the most accurate (PCC and SPCC) imputation without the boundary artifacts or signal over-expression seen in competing methods.

### Integration of FOV-induced incomplete spatial multi-omics data from mouse embryonic brain

In this section, we performed experiments on a mouse embryonic mouse brain dataset spanning the E13.5, E15.5, and E18.5 development stage. Each section was sequenced using MISAR-seq technique, represented by spatial gene expression (RNA) and chromatin accessibility (ATAC) (Figure 3g). To simulate FOV-induced incompleteness, a contiguous 50% region of the RNA modality was *in silico* masked (Figure 3h).

Figure 3i illustrates the domain identification results of PRISM as well as others on the E13.5 brain section. The baseline methods struggled to accurately identify key brain regions, such as the dorsolateral pallium ventral part (DPallv) and dorsolateral pallium medial part (DPallm), which are crucial for understanding distal patterning and neuronal migration. This could be due to incomplete gene expression patterns, which lead to noise interference in these regions. In contrast, PRISM can accurately identify and maintain spatial continuity, as its mechanism also effectively reconstructs relationships between the ATAC and RNA data. Quantitative evaluations of multiple indicators further confirmed the robust performance of PRISM, with its ARI and NMI exceeding those of the second-best baseline (SpaMosaic) by 15.35% and 18.01%, respectively (Figure 3j). Consistently superior performance was observed across all other developmental timepoints (Supplementary Figure 7a-c and 8a-c). Moreover, under fully registered ATAC and RNA inputs, PRISM continued to display exceptional efficacy (Supplementary Figure 9b-d).

Imputation performance was rigorously evaluated (using PCC and SPCC) on the top 800 highly variable genes (HVGs), which represent the highly dynamic features in the dataset. PRISM’s PCC and SPCC correlation distributions for this task exhibited the highest median and the lowest proportion of low-correlation outliers (Figure 3k and Supplementary Figure 9e). These results highlight PRISM’s capability to robustly impute this critical feature set. Other qualitative imputation results can be found in Supplementary Figure 7e-f and 8e-f, further highlighting its advancements.

### Integration of resolution-induced incomplete spatial multi-omics data from PD striatal human brain

PD is a progressive neurodegenerative disorder characterized by motor deficits. Here, we utilized a PD striatal human brain dataset generated via SMA technology (coupling SRT and MALDI-MSI), consisting of three spatially resolved sections, i.e., Cd1, Cd2, and Cd3 in Figure 4b. Although the spatial transcriptomics and metabolomics data originated from the same tissue section, they lacked inherent co-localization due to significant disparities in resolution and coordinate systems, precluding direct multi-omics analysis. We applied the MAGPIE registration tool with a rigorous filtering protocol (Figure 4a), identifying spatial spots with less than 10% overlap and masking the metabolomics values of these low-confidence regions. (Figure 4c and Supplementary Figure 10a). This process, which affected approximately 50% of spatial spots (including mounting artifact regions), produced realistically resolution-induced incomplete spatial multi-omics data.

Figure 4d, e illustrate the domain identification results across three sections with manual annotated ground truth. Qualitative assessment revealed that while PRISM and COSMOS generally outperformed other baselines in delineating spatial domains, only PRISM successfully resolved the complex dopamine distribution pattern within the Caudate (Cd) region (Figure 4e). This capability is essential for distinguishing basal ganglia function from PD-related pathology. In section Cd3, PRISM accurately reconstructed boundaries in sub-regions lacking metabolite measurements, maintaining the continuity of surrounding spatial signals. Notably, the mounting artifact region was predominantly assigned to ‘Cluster 1’, forming distinct boundaries with adjacent functional domains (Figure 4g). Differential expression analysis confirmed the biological validity of this classification (Figure 4h): expression patterns of synaptic markers (e.g., *SYNPR, SCG2, RAB3C, THY1*) and neuronal activity genes (e.g., *FAM171B, CHN1*) within the artifact region were highly consistent with ‘Cluster 1’, yet significantly distinct from other clusters. Quantitative evaluation using established clustering metrics demonstrated that PRISM achieved the best performance across all indices (Figure 4f and Supplementary Figure 10c).

We next assessed the imputation performance by investigating the spatial reconstruction of three specific metabolites directly linked to the dopaminergic response, i.e., *m/z 675*.*28296, m/z 674*.*28592*, and *m/z 691*.*27831* in Supplementary Figure 10d. PRISM successfully recovered signals obscured or distorted within the artifact regions, restoring intensity ranges that closely recapitulated the raw, unfiltered measurements. In contrast, single-cell methods (e.g., cTP-net, scArches, and sciPENN) exhibited distinct artificial stripe-like discontinuities, whereas the spatial baseline SpaMosaic, despite smoothing these artifacts, suffered from signal over-expression in certain sections.

Collectively, these findings highlight PRISM’s exceptional robustness against resolution-induced incompleteness, validating it as a solid framework for reconciling incomplete spatial multi-omics data.

### Integration of incomplete spatial multi-omics data from adjacent sections P22 mouse brain

In this section, we utilized a P22 mouse brain dataset generated via spatial CUT&Tag-RNA-seq. We selected two anatomically adjacent coronal sections to serve as distinct data sources for the spatial epigenome and transcriptome, respectively. Due to inevitable tissue distortion between sections, the data lacked inherent cell-to-cell spatial correspondence, still precluding direct joint multi-omics analysis. Here, we applied the SLAT alignment tool with a rigorous filtering protocol (Figure 5a), discarding cells with alignment similarity scores below 0.65 (Figure 5b, c). This procedure yielded an incomplete spatial multi-omics dataset, comprising intact spatial epigenomic data alongside approximately 50% of the randomly registered transcriptomic data (Figure 5d).

**Fig. 5:**
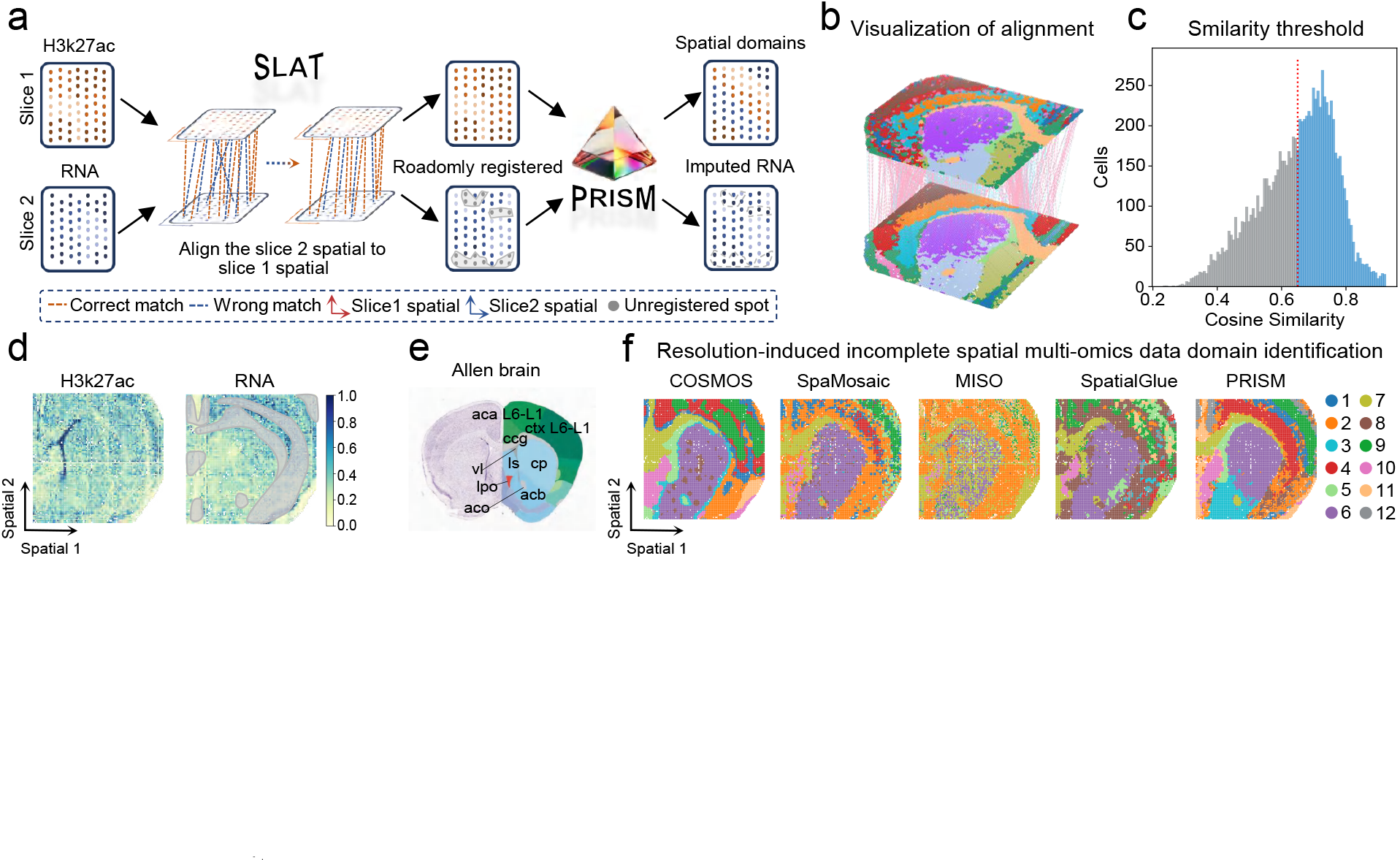
Evaluation of PRISM under simultaneous inconsistent FOV and resolution in adjacent sections P22 mouse brain. **a**, Pipeline schematic: H3K27ac (section 1) and RNA (section 2) from adjacent sections are aligned using SLAT to reconcile morphological distortions. Low-confidence matches are filtered, yielding incomplete spatial multi-omics data characterized by simultaneous inconsistent FOV and resolution. PRISM then infers spatial domains and imputes RNA in unregistered locations. **b**, Visualization of the adjacent sections alignment process, showing correspondences between cells across the two sections after transformation. **c**, Distribution of cosine similarity scores for candidate adjacent sections matches. The red dashed line indicates the threshold (0.65) used to exclude low-confidence matches, resulting in the final incomplete spatial multi-omics dataset. **d**, Spatial signal maps of H3K27ac (section 1) and RNA (section 2) after preprocessing, with the unregistered region indicated. **e**, Reference anatomical annotations from the Allen Brain Atlas for the corresponding coronal section. **f**, Visual comparison of spatial domains identified by PRISM versus state-of-the-art baselines.

Figure 5e, f illustrate the spatial domain identification results benchmarked against the Allen Brain Atlas. Compared with other baseline methods (e.g., SpaMosaic, MISO), PRISM and COSMOS better preserved the spatial continuity of tissue architecture (Figure 5f). However, COSMOS still exhibited fine-scale inaccuracies similar to the other baselines. These limitations were manifested as spatial discontinuities across cortical layers and the erroneous conflation of the caudate putamen (CP) and the nucleus accumbens (ACB) distinct nuclei governing reward processing and motor control. This misclassification likely arises from the loss of transcriptomic heterogeneity necessary to resolve these fine structures during the registration process. By contrast, PRISM accurately separates these functional regions and produces spatial domain boundaries that are most consistent with the reference brain atlas.

We visualized the imputation of key neuronal markers, including *Atp1b1, Kif1a, Snhg11*, and *Syt1* (Supplementary Figure 11). The results demonstrated PRISM’s capability to accurately reconstruct gene expression even in registration-filtered adjacent sections. Notably, the laminar gradient of *Syt1* and the striatal enrichment of *Pde10a* were restored with high fidelity, closely mirroring the expression patterns observed in the target section.

These results indicate that PRISM’s working mechanism effectively mitigates inherent errors stemming from incompatible acquisition protocols, and its compatibility with alignment tools enables analyses comparable to precise registration.

### 3. DISCUSSION

We developed PRISM, a niche-informed adaptive learning framework for spatial multi-omics under incomplete registration. By conceptualizing each cell or spot within its local neighborhood as a coherent “niche”^64,65^, PRISM propagates information along inter-niche similarity maps from registered to unregistered regions and jointly performs spatial domain identification and spatial omics imputation within a unified model. Across various FOV-induced and resolution-induced incompleteness, PRISM consistently outperformed existing methods, demonstrating exceptional utility in reconciling fragmented data and seamless compatibility with established registration pipelines.

Using spatial multi-omics datasets generated by diverse technologies (e.g., MISAR-seq, and SMA), PRISM integrates cross-modal evidence to recover smooth expression gradients and coherent spatial domains. Its performance is comparable to the results obtained under precisely registered conditions. By leveraging inter-niche similarity propagation inspired by COVET, the model captures cross-modal correspondences at the level of cellular niches. For example, the framework can capture the co-distribution of dopamine metabolism-related gene programs and metabolic features in the PD striatum. It uses these relationships as biologically grounded priors to guide the imputation process. Consequently, PRISM restores tissue patterns that are often weakened or lost under single-modality or strictly pointwise registered analyses. It produces spatial domains that better reflect underlying organization, including the reintegration of mounting-artifact regions into clusters that are consistent with dopaminergic annotations in the PD datasets.

While PRISM demonstrates robust capabilities, several avenues warrant further exploration. The field is currently witnessing the rise of large-scale spatial atlases and foundational models that encode rich regulatory semantics^66,67^. Incorporating such pretrained representations into PRISM may further improve the accuracy of cross-modal integration and imputation. At the same time, spatial multi-omics technologies are moving toward higher resolution and broader coverage. Multiple modalities are being measured at the same physical site and across micro to macro scale organization. Adapting PRISM to explicitly handle these high-dimensional and multi-scale settings is an important direction for future work. This may eventually involve more than two omics layers and the integration of hierarchical spatial structures.

In summary, PRISM offers a novel framework that integrates incomplete spatial multi-omics data as these technologies are increasingly used to dissect complex biological processes. It successfully imputes spatial omics and recovers high-quality spatial domains under both FOV-induced and resolution-induced incompleteness, ensuring accurate downstream biological interpretation.

## METHODS

### Data

#### Spatial transcriptomics and proteomics data of human lymphoid organs

Human lymphoid organs, particularly lymph nodes and tonsils, have complex tissue architectures that are central to host immune defense. To profile these tissues *in situ* across molecular layers, the Spatial Transcriptomics-Proteomics (STP) workflow^68^ is designed to simultaneously capture spatially resolved transcript and protein expression, while preserving the specificity and sensitivity of both modalities. The pipeline includes tissue sectioning, spatial transcriptomic profiling, immunofluorescence staining, laser capture microdissection, and spatial proteomics analysis.

In this study, STP datasets from human lymph nodes and tonsils were internally generated using the SpaMosaic framework, yielding fully co-registered spatial transcriptomic and proteomic measurements with expert-derived domain annotations. While SpaMosaic originally focused on a mosaic-style imputation setting, it did not explicitly evaluate scenarios in which the two modalities are available on incomplete spatial multi-omics data. We next modeled this scenario by preserving transcriptomic profiles for all spatial spots in each section while progressively restricting the proteomic modality via masking a subset of spots. Concretely, proteomic masking was implemented in two steps based on cellular spatial coordinates: (i) defining either a horizontally (or vertically) contiguous window that simulates a reduced FOV, or a randomly sampled subset of spots, and (ii) setting protein intensities to zero outside the selected window. This procedure produced sections with full FOV spatial transcriptomics but only partially overlapping proteomics, with effective multi-omics overlap controlled at 90%, 70%, 50%, 30%, and 10%, defined as the fraction of spots retaining registered RNA-protein measurements in each scenario. The same strategy was applied to both tonsil and lymph node tissue sections (Figure 2c and 3b).

#### Microfluidic indexing-based spatial assay for ATAC and RNA-sequencing embryonic mouse brain data

The mouse embryonic brain spatial multi-omics data were obtained from the public repository OEP003285 (https://www.biosino.org/node/project/detail/OEP003285). The dataset was generated by MISAR-seq^8^ and covers three fully registered developmental stage sections (E13.5, E15.5, E18.5) with annotation of functionally distinct brain regions. Each section contains 191,034 ATAC peaks and 24,368 gene-level RNA measurements. We next simulated a FOV-induced incompleteness scenario, in which the RNA modality was available only on a reduced FOV relative to ATAC. Specifically, we focused on the 3,000 HVGs, which captured the majority of informative transcriptional variations, and set their expression values to zero within 50% of a spatially contiguous region (Figure 3h). This procedure yielded sections in which ATAC measurements remained available across the full FOV, whereas RNA measurements were retained only within the remaining half of the spatially contiguous region. The same masking strategy was applied to all three developmental-stage sections.

#### Spatial transcriptomic-metabolomics PD striatum human brain data

The SMA workflow was applied to three coronal striatal sections at distinct positions, denoted Cd1, Cd2, Cd3, obtained from a frozen postmortem human PD striatal tissue. Each section provided spatial transcriptomics (generated with 10x Genomics Visium), H&E histology, spatial metabolomics (measured by MALDI-MSI), and expert annotations of brain regions and dopamine distribution. We used these annotations to aggregate spatial spots into brain-region level dopamine response labels, which then served as ground truth for downstream analysis and evaluation (Figure 4d).

Due to technical constraints, including inconsistent spatial resolution, sequential (non-simultaneous) assays, and mounting artifacts, the transcriptomic and metabolomic layers were not fully co-registered. We refined the registration using MAGPIE, leveraging the H&E image as a common coordinate system to map Visium spots and MALDI-MSI pixels into a shared space (Figure 4a). The procedure comprised four main steps: (i) landmark selection, (ii) coordinate transformation and co-registration, (iii) construction of a Space Ranger-style Visium object, and (iv) cross-modal matching and aggregation. We retained only spots with sufficient cross-modal overlap as valid registered locations. For all other spatial spots, including those within mounting-artifact regions, metabolite intensities were set to zero while RNA measurements were preserved at all Visium locations (Supplementary Figure 10a, b).

#### Spatial histone modifiemics-transcriptomics P22 mouse brain data

In the P22 mouse brain datasets, spatial histone modifications and transcriptomes were jointly profiled at near single-cell resolution using CUT&Tag-RNA sequencing. This dataset two anatomically adjacent coronal sections were selected and used one as the source of the spatial histone-modification map and the other as the source of the spatial transcriptomic map. We then used SLAT to perform cell-level registration between the two sections (Figure 5a), correcting inherent biases between adjacent sections, such as tissue deformation and the lack of one-to-one cell-level spatial correspondence. The SLAT workflow comprises five steps: (i) preprocessing and cell/spot localization, (ii) construction of morphological and molecular features, (iii) similarity estimation and candidate pairing, (iv) coordinate transformation and alignment refinement, and (v) threshold-based selection of matched pairs. This process excluded 4,886 cells (approximately 50%) with similar scores below 0.65 from the aligned space and set their gene expression values to zero (Figure 5c). This SLAT-based workflow produced near single-cell incomplete spatial multi-omics data, exemplifying the challenges stemming from the alignment of distorted adjacent sections.

### PRISM for incomplete spatial multi-omics data

PRISM is a niche-informed spatial multi-omics framework designed to operate under conditions of incomplete registration. It jointly imputes the spatial omics and delineates spatial domains, thereby reconstructing tissue-level continuity and yielding a coherent multi-omics atlas. The approach is robust across both FOV-induced and resolution-induced incompleteness scenarios and interoperates with existing registration pipelines, delivering performance that closely matches analyses based on fully registered spatial multi-omics data.

For each cell or spatial spot with multi-omics measurements, PRISM empolyed two graph-attention auto-encoders^45^ to learn representations of the source and target modalities, respectively. Each encoder consisted of two graph-attention convolutional layers followed by linear transformations, capturing both node attributes and spatial relationships to represent complex expression patterns. Methodologically, PRISM was inspired by COVET. It leveraged a inter-niche similarity map to identify the top-*k* most similar neighbours for each unregistered cell or spatial spot and used their averaged expression profile as an initial estimate of the target modality. After the initialization, source and target omics features were concatenated and fed into Transformer encoder to model long-range cross-modal dependencies and to refine the target-modality representation. Two subsequent linear blocks (composed of fully connected layers with ReLU activation) projected the Transformer output into source and target specific subspaces, yielding modality-specific embeddings. A feature interaction module then integrated the source embedding, target embedding, and cross-modal relations into a unified semantic representation for spatial domain identification. Finally, two omics-specific decoders (composed of two graph-attention convolution layers and inverse linear transformations) for the source and target omics were performed separate tasks: reconstructing the source omics representation and predicting the target omics data to be registered.

### Data preprocessing

For a section containing *n* cell or spatial spot, the original expression matrix of source omics *S* and target omics *T* are defined as 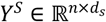 and 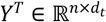, respectively. While both modalities are sampled from the same *n* spatial locations, *Y*^*T*^ represents an incomplete target profile where a subset of cells/spots lacks measured expression counts, a condition referred to as partial pairing. To formally represent this incompleteness, we define a binary mask matrix ℳ_*i*_ ∈ {0,1}^*n* ×1^. when ℳ_*i*_ = 1 indicates a registered cell/spot with complete spatial multi-omics measurements, and ℳ_*i*_ = 0 marks an unregistered cell/spot where the target omics is missing the original expression counts. Here, *U* = *S* ∪ *T* ∈ {*m*_1_, …, *m*_*k*_} denotes the union of all source and target omics, including spatial histone modifications (epigenomics), chromatin accessibility, transcriptomics, proteomics, and metabolomics. For each modality *m*_*i*_, we applied modality-specific preprocessing: (i) spatial transcriptomics: genes detected in fewer than 10 cells were removed, followed by library-size normalization and log1p transformation, (ii) spatial epigenomics and chromatin accessibility: term frequency-inverse document frequency transformation was applied, and (iii) proteomics and metabolomics: intensity normalization and logarithmic transformation were performed. After preprocessing, we independently selected the top 3,000 highly variable genes, the top 50,000 highly variable epigenomic features, and the top 100 highly variable metabolites, and retained all protein features. These biologically representative spatial multi-omics expressions (i.e., 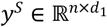 and 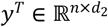) are used as the input for the subsequent encoder.

### Construction of COVET-based similarity prior

Compared to single-modality spatial omics, spatial multi-omics integrates information across omics layers, which is crucial for deciphering complex biological states of cells and tissues. To bridge cross-modal associations for each cell/spot, PRISM extracted the inter-niche similarity covariance covariation patterns based on the COVET prior. During niche construction, each cell/spot *i* formed a niche 𝒩_*i*_ by identifying its spatial neighbors via the k-nearest neighbors (kNN) algorithm:

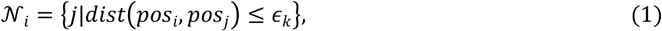

where *k* was set as 6 for spot-resolution datasets and 8 for cell-resolution datasets. Inspired by COVET, PRISM defined niche characteristics also using “Shifted Covariance” and used the average of entire section 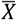 as a reference, ensuring comparability across niches:

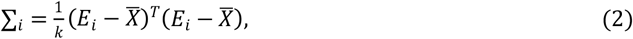

Where 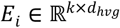 represents the expression matrix within the neighborhoods. The inter-niche similarity is then quantified by approximate optimal transport (AOT)^43,69^ distance. According to the derivation by COVET, when the covariance matrix is a positive definite matrix, the AOT distance can be equivalent to the squared euclidean distance between the matrix square root of the covariance matrix:

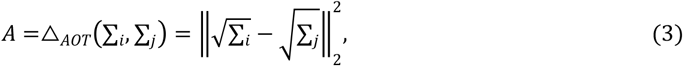

This transformation avoids the complex matrix multiplication, significantly enhancing the computational efficiency on large-scale datasets. For each cell/spot, all other niches are reordered based on their AOT distances from closest to farthest, identifying the most similar cells/spots. The inter-niche similarity matrix *A* ∈ ℝ^*n*×*n*^ is then constructed to guide PRISM modeling.

### Graph-Attention encoding and niche retrieval

The preprocessed matrices *X*^*m*^, where *m* ∈ {*S, T*}, and the spatial adjacency matrix *P* are fed into two layers Graph Attention (GAT) encoders. By capturing both individual cell/spot attributes and their local spatial relationships, the encoders map the features into a low-dimensional latent space:

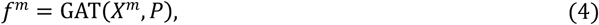

where *f*^*s*^ and *f*^*T*^ represent the encoded features for the source and target omics, respectively. For each cell/spot *i*, PRISM reorders all other cells/spots based on their AOT distances to retrieve the top-*k* most similar registered cells/spots (ℳ_*k*_ = 1), denoted as the set 𝒩_*retrieved*_ ∈ ℝ ^*n*×*k*^. The enhanced source representation 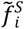 and initial target estimate 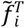 are concurrently generated by aggregating the encoded features from the same set of retrieved registered neighbors.

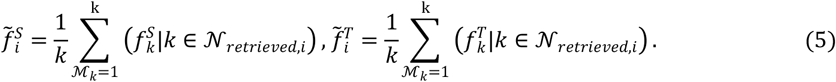

This niche-informed symmetric projection imbues both modalities with shared context to maintain tissue-level biological continuity in unregistered regions while ensuring synchronized cross-modal refinement in the subsequent Transformer encoder stage.

### Similarity-guided imputation tokens

For incomplete registered spatial multi-omics data, we enriched the source and target omics context by integrating inter-niche similarity, while maximizing recovery of missing target features. Specifically, learnable tokens were introduced to label cells/spots requiring imputation (ℳ_*i*_ = 0). These tokens enable the model to infer enrichment features from the top-*k* similar neighbors in registered regions while approximating target profiles for unregistered cells/spots. The integration of these tokens with aggregated features 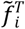 establishes contextual embeddings between cells/spots and their similar counterparts. All tokens were initialized from *N*(0,1) for symmetric feature propagation. This configuration coordinates with subsequent multi-task loss functions to ensure high performance while preventing the cascading error accumulation inherent in sequential processing.

### Transformer encoder

Transformer architecture has been widely adopted in single-cell omics and spatial omics analysis^66,67,70,71^. In PRISM, the Transformer encoder module established inter-omics communication, further recovering unmeasured omics features of unregistered cells/spots. It comprised two core components, multi-head self-attention and feed-forward networks, linked by residual connections and layer normalization to improve training stability and representation capacity.

As illustrated in Figure 1d, the Transformer’s input feature *F*_*i*_ is concatenated by the enhanced source representation 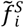 and the target representation 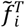. For efficient multi-head attention processing, these concatenated features are distributed into *c* batches, with each batch feature denoted as *F*_*i*_ ∈ [*F*_1_, …, *F*_c_].The module took query (*Q*), key (*K*), and value (*V*) vectors as inputs to compute inter-position correlations. The computational workflow was:

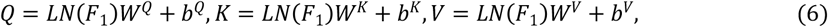

where *LN* represents layer normalization, and *W*^*Q,k,V*^ and *b*^*Q,K,V*^ represent learnable transformation matrix and bias, respectively. Following the multi-head strategy, we split the features dimension of size *m* into *d* segments, with each segment corresponding to independent set of *Q,K,V* ∈ ℝ^*N*×*d*×(*m*/*d*)^. These segmented features were then processed in parallel in multiple subspaces using scaled dot-product attention followed by softmax normalization, yielding the attention weights for each head. For a single head (*h* = 1), the attention computation reduces to:

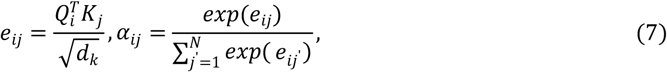

where *i* and *j* are the position indices of *Q* and *K*, respectively, and 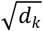 is the scaling factor that normalizes the dot product by the key dimension. Each element *α*_*ij*_ in the attention matrix indirectly reflects the degree of communication between cells during the target omics recovery process. The attention output of a single head is obtained by a weighted sum of value vectors such as:

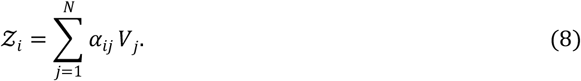

For *H* heads, we concatenated the head-wise outputs and applied a linear projection to obtain the multi-head attention output:

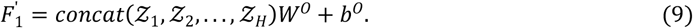

We then added a residual connection from the input 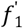 and applied layer normalization as:

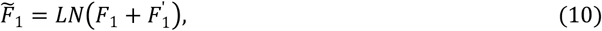

and use 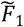 as the input to a position-wise FNN. The FNN performed deep feature recalibration through two linear layers with ReLU activation and a second residual connection:

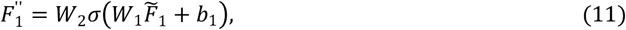

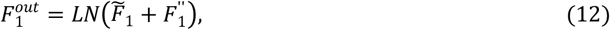

where *σ*(·) denotes the ReLU function. The final output of this Transformer encoder over all batch can be expressed as 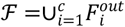.

Cellular phenotypes in tissues are regulated by both intrinsic gene expression and niche interactions ^71^. This registers with our core hypothesis: cells or spatial spots with analogous spatial contexts are likely to display similar expression profiles. Transformer architecture in PRISM is designed to effectively infer and restore the omics information of cells in unregistered regions, while fully utilizing biological prior knowledge and context-rich spatial multi-omics information.

### Semantic embedding

While single-modality spatial omics suffices for certain scenarios, multi-omics integration becomes essential as cellular phenotypes emerge from cross-modal interactions^11,51,72^. This inter-omics regulation constitutes a cornerstone of our hypothesis. For instance, STP analysis requires modeling mRNA-protein expression relationships to elucidate translational control and protein stability mechanisms. Thus, deriving integrated semantic embeddings from spatial multi-omics features is critical. The attention features were processed through a dual-linear block (ReLU activation), then projected into distinct feature spaces (ℱ^*S*^ and ℱ^*T*^) via source- and target-decoder pathways. Feature interaction operations then generated advanced cross-modal representations^36^. For source and target omics, the interaction matrix for spot/cell *i* was computed as:

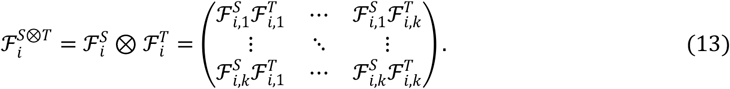

For each omics, all spots interaction matrices were flattened, then concatenated into a spot-level interaction feature matrix. PCA was performed on each matrix to extract the top-*k* principal components, which were integrated as complementary signals into the unified embedding space. This enhanced downstream analysis compatibility and connected selected interaction features with pattern-specific feature vectors, ultimately forming the integrated semantic embeddings of PRISM.

### Feature decoder

PRISM incorporates four modality-specific decoders, each structured with two graph-attention convolution layers followed by inverse linear transformations. This spatial-aware configuration leverages local neighborhood topologies to facilitate the precise recovery of niche-specific molecular distributions and global tissue patterns. The decoding process operates through two distinct functional pathways. The first pathway is decoders 1 and 3 prioritize data fidelity by mapping the initial GAT-encoded embeddings (i.e., *f*^*s*^ and *f*^*T*^) back to their original feature spaces. The resulting reconstructed profiles for the source and target modalities are denoted as 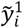 and 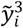, respectively. Simultaneously, the another one is that decoders 2 and 4 leverage the projected representations (i.e., ℱ^*s*^ and ℱ^*T*^) to produce enhanced source profiles denoted as 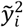 and final predicted target profiles denoted as 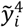 for unregistered regions. To maintain biological consistency and parameter efficiency, modality-specific weight sharing (e.g., decoders 1 and 2 as well as 3 and 4) is implemented between the respective reconstruction and refinement decoders. This integrated framework enables robust source preservation while facilitating accurate cross-modal imputation.

### Multi-task objective function and optimization

PRISM optimization leverages a multi-task objective function grounded in Mean Squared Error (MSE) to synchronize spatial omics reconstruction and imputation. This multi-stage objective function is strictly governed by the asymmetric spatial availability of reference data. While the source modality ground-truth 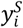 is provided across all *n* cells or spatial spots, the target-modality ground-truth 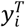 is observed exclusively at registered spots where ℳ_*i*_ = 1. This data sparsity necessitates a dual-stage supervision strategy where the model first constrains initial embeddings to maintain data fidelity and then supervises refined embeddings to facilitate deep inter-modality communication and molecular manifold restoration. This joint optimization establishes a reciprocal dependency between domain context and molecular manifolds, avoiding cascading errors through end-to-end gradient synchronization. The loss functions are formally expressed as follows:

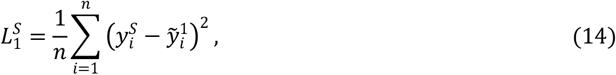

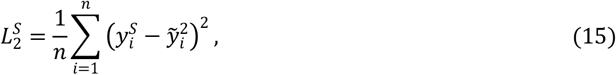

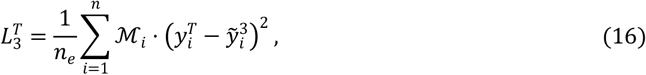

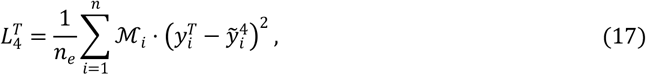

Where *n* and *n*_*e*_ denote the total and registered spot counts, 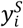 and 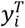 denote the ground-truth expression counts for the source and target modalities at spot *i*, respectively. Here, 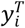 is only available exclusively for registered cells/spots (i. e., ℳ_*i*_ = 1), since there is no ground-truth expression counts for the target omics at the unregistered cells/spots.

Dynamic balancing of these diverse loss components is achieved through learnable normalized weights *β*_*i*_ ∈{ *β*_1_, …, *β*_*n*_}, which harmonize the reconstruction and imputation pathways during training. These weights are derived using an exponential formula to ensure their sum remains equal to one:

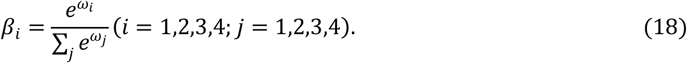

Finally, the total loss function integrates these weighted contributions to guide the update of the entire model parameters:

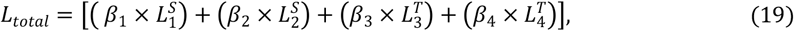

Where ∑ *β*_*i*_ = 1, and *ω*_*i*_ was the initialized weighting coefficient.

### Evaluation criteria

The regression and multi-class classification tasks in this study were evaluated using distinct metrics. For the regression task, both PCC and SPCC were employed as correlation evaluation criteria. PCC measures linear correlation, with its formula presented below:

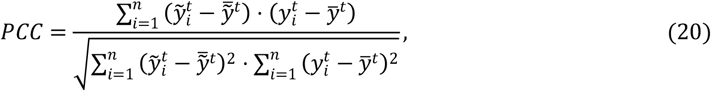

where 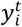 and 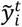 represent the ground truth and transformed values respectively, with 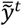 and 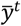 denoting their mean values. SPCC primarily measures monotonic relationships, formulated as:

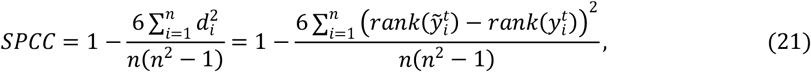

where *n* is the sample size, and *d*_*i*_ denotes the rank difference between ground truth and predicted values for the cell/spot *i*. For the spatial domain recognition multi-classification task, we employed AMI, NMI, ARI, Homogeneity, and V-measure. These evaluation metrics are formally defined as follows:

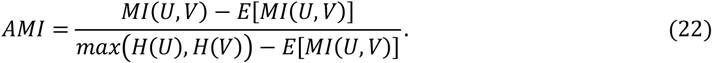

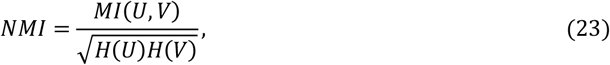

where *M I*(*U,V*) represents the mutual information, *H*(*U*) and *H*(*V*)denote the entropy of clustering results U and V respectively, and *E*[*MI*(*U,V*)] indicates the expected value adjusted for chance effects.

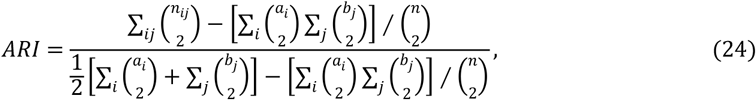

where *n* is the total number of objects in the dataset, *a*_*i*_ denotes the number of objects in the *i*^*th*^ cluster of the ground truth partition, *b*_*j*_ represents the number of objects in the *j*^*th*^ cluster of the algorithmic partition, *n*_*ij*_ indicates the number of objects belonging to both the *i*^*th*^ and *j*^*th*^clusters.

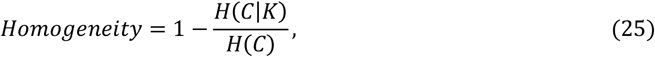

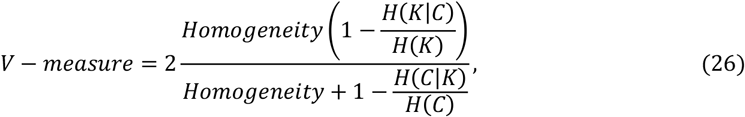

where *H*(*C*|*K*) represents the conditional entropy, *H*(*C*) denotes the entropy of ground truth classes *C*, while *H*(*K*|*C*) indicates the conditional entropy, and *H*(*K*)measures the entropy of clustering results *K*.The model was evaluated using the metrics, where higher values consistently indicate better clustering performance across all metrics. For datasets comprising multiple tissue sections, standardized evaluation criteria were uniformly applied with corresponding results visualized.

### Simulated datasets for incomplete scenario analysis

We systematically evaluated PRISM’s performance under FOV-induced incompleteness using three simulation datasets derived from fully registered spatial multi-omics data. These datasets covered spot-resolution human lymphoid organs and mouse brains and spanned three modality combinations: RNA-protein, ATAC-RNA, and RNA-metabolomics. The human lymphoid organ collection comprised two datasets (lymph node and tonsil), each consisting of three tissue sections, with each section containing expression profiles for up to 18,085 genes and 31 proteins. The mouse brain dataset captured embryonic brains at three developmental stages and included chromatin accessibility and gene expression data from three consecutive coronal sections, each containing 32,285 genes and 161,461 chromatin peaks.

### Real datasets for incomplete scenario analysis

We assessed PRISM’s performance on real-world incompatible platforms and its compatibility with existing alignment tools using two incomplete spatial multi-omics datasets, spanning resolution-induced incompleteness on a single section and simultaneous inconsistent FOV and resolution across adjacent sections. The first dataset was a human PD striatum dataset comprising three coronal sections, each with RNA (Visium) and metabolomics (MALDI-MSI). Cross-modal registration was performed with a MAGPIE-based pipeline, yielding a spot-resolution spatial multi-omics dataset characterized by resolution-induced incompleteness. The second dataset was a P22 mouse brain data consisting of two anatomically adjacent coronal sections, each profiled for H3k27ac and RNA at near cell-resolution. Adjacent sections registration was performed with a SLAT-based pipeline, yielding a near single-cell incomplete spatial multi-omics data characterized by simultaneous inconsistent FOV and resolution.

### Implementation details

This study was implemented on the PyTorch platform. Models were trained using the Adam optimizer with learning rate=1e-3 and weight decay=1e-4. To optimize computational resource utilization, after 200 training epochs, we employed early stopping with patience=20, terminating training if no improvement occurs for 20 consecutive epochs. The maximum epoch limit was set to 1,000, with a batch size of 64. All experiments ran on a single NVIDIA A100 40GB graphics card, ensuring efficient computation for spatial multi-omics data.

## Author contributions

S.M., Z.W. and Y.L. contributed equally to this work. Z.Y. and C.H. jointly supervised the project. S.M., Z.W. and Y.L. conceived the study, developed and implemented the computational methodology, performed the experiments and data analyses, and wrote the original draft. Z.Y. provided guidance on study design and experimental validation. C.H. supervised the project, provided guidance throughout, and supported the experimental work. J.L. and D.L. assisted with figure refinement and manuscript editing. C.W. and J.X. contributed to figure preparation and manuscript revision. X.S., T.Z. and W.H. contributed to analysis planning and manuscript revision. J.S. provided suggestions for the work and guidance on manuscript revision. All authors reviewed and approved the final manuscript.

## Acknowledgements

This work was supported in part by the National Key Research and Development Program of China under Grant 2022YFE0112200; in part by the Key Research and Development Program of Guangzhou under Grant 2023B01J1001, Grant 2023B01J0002; in part by the National Natural Science Foundation of China under Grant 62502161, Grant 62325204, Grant U21A20520, Grant 62102153, Grant 62272326, Grant 62502163, and Grant 62172112; in part by the China Postdoctoral Science Foundation (Certificate Number: 2025M782912). in part by the Science and Technology Project of Guangdong Province under Grant 2022A0505050014, Grant 2025A04J5480; in part by the Key-Area Research and Development Program of Guangzhou City under Grant 202206030009; in part by the Natural Science Foundation of Guangdong Province of China under Grant 2022A1515011162 and Grant 2023A1515012894; in part by the Guangdong Natural Science Funds for Distinguished Young Scholar under Grant 2023B1515020097.

## Inclusion & Ethics

Not relevant.

## References

1 Chen, J. G. et al. Giotto Suite: a multiscale and technology-agnostic spatial multiomics analysis ecosystem. Nature Methods 22, 2052–2064 (2025).

2 Liu, X. et al. Spatial multi-omics: deciphering technological landscape of integration of multi-omics and its applications. Journal of Hematology & Oncology 17, 72 (2024).

3 Vickovic, S. et al. High-definition spatial transcriptomics for in situ tissue profiling. Nature methods 16, 987–990 (2019).

4 Vandereyken, K., Sifrim, A., Thienpont, B. & Voet, T. Methods and applications for single-cell and spatial multi-omics. Nature Reviews Genetics 24, 494–515 (2023).

5 Liu, Y. et al. High-plex protein and whole transcriptome co-mapping at cellular resolution with spatial CITE-seq. Nature Biotechnology 41, 1405–1409 (2023).

6 Ståhl, P. L. et al. Visualization and analysis of gene expression in tissue sections by spatial transcriptomics. Science 353, 78–82 (2016).

7 Vicari, M. et al. Spatial multimodal analysis of transcriptomes and metabolomes in tissues. Nature Biotechnology 42, 1046–1050 (2024).

8 Jiang, F. et al. Simultaneous profiling of spatial gene expression and chromatin accessibility during mouse brain development. Nature Methods 20, 1048–1057 (2023).

9 Zhang, D. et al. Spatial epigenome–transcriptome co-profiling of mammalian tissues. Nature 616, 113–122 (2023).

10 Hu, J. et al. SpaGCN: Integrating gene expression, spatial location and histology to identify spatial domains and spatially variable genes by graph convolutional network. Nature methods 18, 1342–1351 (2021).

11 Long, Y. et al. Deciphering spatial domains from spatial multi-omics with SpatialGlue. Nature Methods 21, 1658–1667 (2024).

12 Sun, C. et al. Spatially resolved multi-omics highlights cell-specific metabolic remodeling and interactions in gastric cancer. Nature communications 14, 2692 (2023).

13 Chee-Huat, L. E. et al. Transcriptome-scale super-resolved imaging in tissues by rna seqfish+. Nature 568, 235–239 (2019).

14 Dong, Y. et al. Transcriptome analysis of archived tumors by Visium, GeoMx DSP, and Chromium reveals patient heterogeneity. Nature communications 16, 4400 (2025).

15 Liao, S. et al. Integrated spatial transcriptomic and proteomic analysis of fresh frozen tissue based on stereo-seq. Preprint at 10.1101/2023.04.28.538364 (2023).

16 Liu, Y. et al. High-spatial-resolution multi-omics sequencing via deterministic barcoding in tissue. Cell 183, 1665-1681. e1618 (2020).

17 Marco Salas, S. et al. Optimizing Xenium In Situ data utility by quality assessment and best-practice analysis workflows. Nature Methods 22, 813–823 (2025).

18 Stickels, R. R. et al. Highly sensitive spatial transcriptomics at near-cellular resolution with Slide-seqV2. Nature biotechnology 39, 313–319 (2021).

19 Xia, C., Fan, J., Emanuel, G., Hao, J. & Zhuang, X. Spatial transcriptome profiling by MERFISH reveals subcellular RNA compartmentalization and cell cycle-dependent gene expression. Proceedings of the National Academy of Sciences 116, 19490–19499 (2019).

20 Zhang, D. et al. Spatial dynamics of brain development and neuroinflammation. Nature 647, 213–227 (2025).

21 Chen, A. et al. Spatiotemporal transcriptomic atlas of mouse organogenesis using DNA nanoball-patterned arrays. Cell 185, 1777-1792. e1721 (2022).

22 Liao, S. et al. Stereo-cell: Spatial enhanced-resolution single-cell sequencing with high-density DNA nanoball-patterned arrays. Science 389, eadr0475 (2025).

23 Liu, N. et al. standR: spatial transcriptomic analysis for GeoMx DSP data. Nucleic Acids Research 52, e2–e2 (2024).

24 Ren, P. et al. Systematic benchmarking of high-throughput subcellular spatial transcriptomics platforms across human tumors. Nature Communications 16, 9232 (2025).

25 Li, Z. et al. Integrative deep learning of spatial multi-omics with SWITCH. Nature Computational Science 5, 1051–1063 (2025).

26 Biancalani, T. et al. Deep learning and alignment of spatially resolved single-cell transcriptomes with Tangram. Nature methods 18, 1352–1362 (2021).

27 Clifton, K. et al. STalign: Alignment of spatial transcriptomics data using diffeomorphic metric mapping. Nature communications 14, 8123 (2023).

28 Li, H. et al. SANTO: a coarse-to-fine alignment and stitching method for spatial omics. Nature Communications 15, 6048 (2024).

29 Liu, X., Zeira, R. & Raphael, B. J. Partial alignment of multislice spatially resolved transcriptomics data. Genome Research 33, 1124–1132 (2023).

30 Zeira, R., Land, M., Strzalkowski, A. & Raphael, B. J. Alignment and integration of spatial transcriptomics data. Nature Methods 19, 567–575 (2022).

31 Goltsev, Y. et al. Deep profiling of mouse splenic architecture with CODEX multiplexed imaging. Cell 174, 968-981. e915 (2018).

32 Kreutzer, L. et al. Simultaneous metabolite MALDI-MSI, whole exome and transcriptome analysis from formalin-fixed paraffin-embedded tissue sections. Laboratory Investigation 102, 1400–1405 (2022).

33 Hu, Y. et al. Benchmarking algorithms for single-cell multi-omics prediction and integration. Nature Methods 21, 2182–2194 (2024).

34 Li, L., Dong, L., Zhang, H., Xu, D. & Li, Y. spaLLM: enhancing spatial domain analysis in multi-omics data through large language model integration. Briefings in Bioinformatics 26, bbaf304 (2025).

35 Zhou, Y. et al. Cooperative integration of spatially resolved multi-omics data with COSMOS. Nature communications 16, 27 (2025).

36 Coleman, K. et al. Resolving tissue complexity by multimodal spatial omics modeling with MISO. Nature methods 22, 530–538 (2025).

37 Yan, X. et al. Mosaic integration of spatial multi-omics with SpaMosaic. Preprint at 10.1101/2024.10.02.616189 (2024).

38 Deng, Y. et al. Spatial-CUT&Tag: spatially resolved chromatin modification profiling at the cellular level. Science 375, 681–686 (2022).

39 Deng, Y. et al. Spatial profiling of chromatin accessibility in mouse and human tissues. Nature 609, 375–383 (2022).

40 Rao, A., Barkley, D., França, G. S. & Yanai, I. Exploring tissue architecture using spatial transcriptomics. Nature 596, 211–220 (2021).

41 Williams, E. C. et al. Spatially resolved integrative analysis of transcriptomic and metabolomic changes in tissue injury studies. Nature Communications 17, 205 (2026).

42 Xia, C.-R., Cao, Z.-J.Tu, X.-M. & Gao, G. Spatial-linked alignment tool (SLAT) for aligning heterogenous slices. Nature Communications 14, 7236 (2023).

43 Haviv, D. et al. The covariance environment defines cellular niches for spatial inference. Nature Biotechnology 43, 269–280 (2025).

44 Vaswani, A. et al. Attention is all you need. Advances in neural information processing systems (2017).

45 Velickovic, P. et al. Graph attention networks. In International Conference on Learning Representations (ICLR) (2018).

46 Zhou, Z., Ye, C., Wang, J. & Zhang, N. R. Surface protein imputation from single cell transcriptomes by deep neural networks. Nature communications 11, 651 (2020).

47 Lotfollahi, M. et al. Mapping single-cell data to reference atlases by transfer learning. Nature biotechnology 40, 121–130 (2022).

48 Lakkis, J. et al. A multi-use deep learning method for CITE-seq and single-cell RNA-seq data integration with cell surface protein prediction and imputation. Nature machine intelligence 4, 940–952 (2022).

49 Ren, H., Walker, B. L., Cang, Z. & Nie, Q. Identifying multicellular spatiotemporal organization of cells with SpaceFlow. Nature communications 13, 4076 (2022).

50 Yuan, Z. et al. Benchmarking spatial clustering methods with spatially resolved transcriptomics data. Nature Methods 21, 712–722 (2024).

51 Zhu, B. et al. CellLENS enables cross-domain information fusion for enhanced cell population delineation in single-cell spatial omics data. Nature Immunology 26, 963–974 (2025).

52 Hu, Y. et al. Benchmarking clustering, alignment, and integration methods for spatial transcriptomics. Genome Biology 25, 212 (2024).

53 Wess, M. et al. Spatial integration of multi-omics data from serial sections using the novel Multi-Omics Imaging Integration Toolset. GigaScience 14, giaf035 (2025).

54 Stoeckius, M. et al. Simultaneous epitope and transcriptome measurement in single cells. Nature methods 14, 865–868 (2017).

55 Khan, M., Arslanturk, S. & Draghici, S. A comprehensive review of spatial transcriptomics data alignment and integration. Nucleic Acids Research 53, gkaf536 (2025).

56 Bao, F. et al. Integrative spatial analysis of cell morphologies and transcriptional states with MUSE. Nature biotechnology 40, 1200–1209 (2022).

57 Cao, Z.-J. & Gao, G. Multi-omics single-cell data integration and regulatory inference with graph-linked embedding. Nature Biotechnology 40, 1458–1466 (2022).

58 Longo, S. K., Guo, M. G., Ji, A. L. & Khavari, P. A. Integrating single-cell and spatial transcriptomics to elucidate intercellular tissue dynamics. Nature Reviews Genetics 22, 627–644 (2021).

59 Hao, Y. et al. Integrated analysis of multimodal single-cell data. Cell 184, 3573-3587. e3529 (2021).

60 He, Z. et al. Mosaic integration and knowledge transfer of single-cell multimodal data with MIDAS. Nature biotechnology 42, 1594–1605 (2024).

61 Argelaguet, R. et al. MOFA+: a statistical framework for comprehensive integration of multi-modal single-cell data. Genome biology 21, 111 (2020).

62 Cui, H. et al. scGPT: toward building a foundation model for single-cell multi-omics using generative AI. Nature methods 21, 1470–1480 (2024).

63 Gayoso, A. et al. Joint probabilistic modeling of single-cell multi-omic data with totalVI. Nature methods 18, 272–282 (2021).

64 Birk, S. et al. Quantitative characterization of cell niches in spatially resolved omics data. Nature Genetics 57, 897–909 (2025).

65 Yuan, Z. MENDER: fast and scalable tissue structure identification in spatial omics data. Nature Communications 15, 207 (2024).

66 Szałata, A. et al. Transformers in single-cell omics: a review and new perspectives. Nature methods 21, 1430–1443 (2024).

67 Tejada-Lapuerta, A. et al. Nicheformer: a foundation model for single-cell and spatial omics. Nature methods 22, 2525–2538 (2025).

68 Ben-Chetrit, N. et al. Integration of whole transcriptome spatial profiling with protein markers. Nature biotechnology 41, 788–793 (2023).

69 Cang, Z. et al. Screening cell–cell communication in spatial transcriptomics via collective optimal transport. Nature methods 20, 218–228 (2023).

70 Hao, M. et al. Large-scale foundation model on single-cell transcriptomics. Nature methods 21, 1481–1491 (2024).

71 Wang, Z. et al. NicheTrans: spatial-aware cross-omics translation. Preprint at 10.1101/2024.12.05.626986 (2024).

72 Dong, K. & Zhang, S. Deciphering spatial domains from spatially resolved transcriptomics with an adaptive graph attention auto-encoder. Nature communications 13, 1739 (2022).

