## Supplemental Figure 1-11 for "PRISM: Niche-informed Deciphering of Incomplete Spatial Multi-Omics Data"

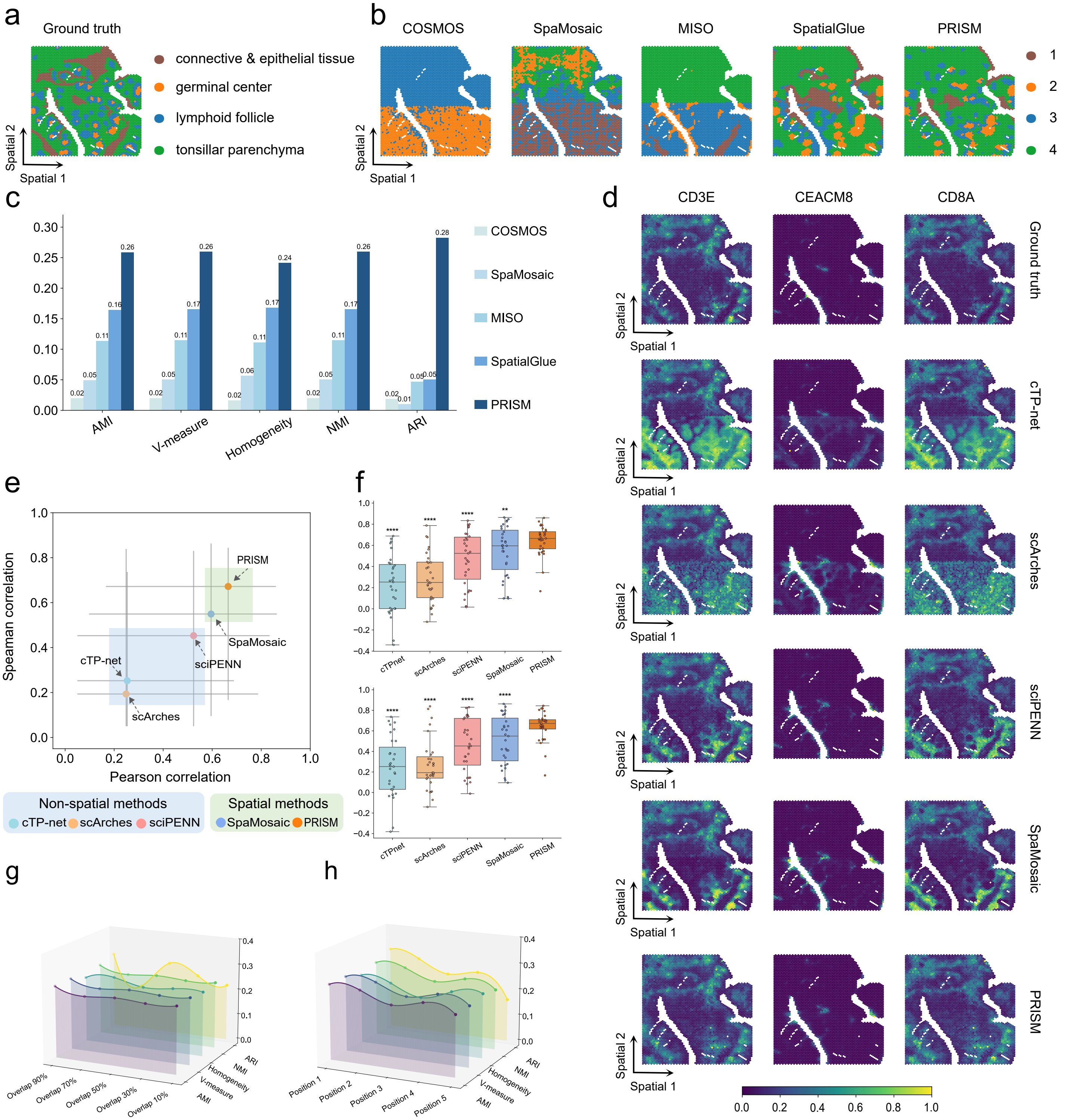


**Supplementary Fig. 1 | Performance of PRISM on human tonsil (slice1) spatial multi-omics under FOV-induced incompleteness.** a, Ground-truth spatial domain annotations for the human tonsil dataset. b, Visual comparison of spatial domains identified by PRISM versus state-of-the-art baselines (COSMOS, SpaMosaic, MISO, and SpatialGlue) under the incomplete spatial multi-omics setting. c,Quantitative benchmarking of domain identification performance using AMI, V-measure, homogeneity, NMI, and ARI. d, Spatial visualization of representative proteins (CD3E, CEACAM8, and CD8A): ground truth measurements compared with imputations from non-spatial translators (cTP-net, scArches, sciPENN) and spatial methods (SpaMosaic, PRISM). e , Scatter plot summarizing overall protein imputation performance (PCC vs. SPCC). Shaded regions distinguish non-spatial methods (blue) from spatial methods (green). f, Box plots illustrating the distribution of protein-specific imputation accuracy (PCC and SPCC). Statistical significance was determined using a two-sided Wilcoxon signed-rank test: *P<0.05, **P<0.01, ***P< 0.001, ****P<0.0001. Center lines indicate medians. g, Robustness analysis of domain identification metrics under varying FOV overlap rates (from 90% down to 10%). h, Robustness analysis regarding the spatial position of the paired window (evaluated across five distinct positions).

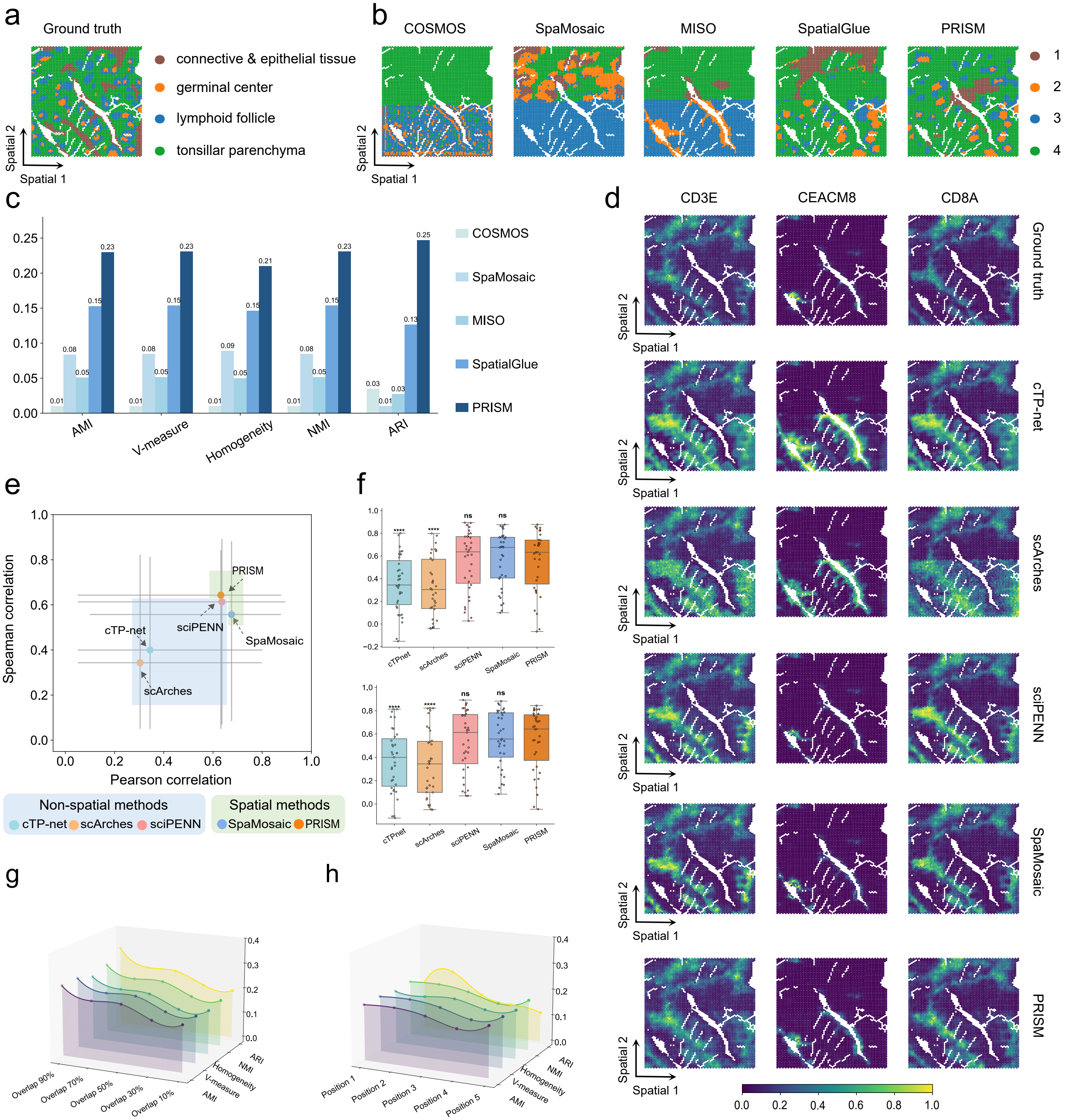


**Supplementary Fig. 2 | Performance of PRISM on human tonsil (slice3) spatial multi-omics under FOV-induced incompleteness.** a, Ground-truth spatial domain annotations for the human tonsil dataset. b, Visual comparison of spatial domains identified by PRISM versus state-of-the-art baselines (COSMOS, SpaMosaic, MISO, and SpatialGlue) under the incomplete spatial multi-omics setting. c,Quantitative benchmarking of domain identification performance using AMI, V-measure, homogeneity, NMI, and ARI. d, Spatial visualization of representative proteins (CD3E, CEACAM8, and CD8A): ground truth measurements compared with imputations from non-spatial translators (cTP-net, scArches, sciPENN) and spatial methods (SpaMosaic, PRISM). e, Scatter plot summarizing overall protein imputation performance (PCC vs. SPCC). Shaded regions distinguish non-spatial methods (blue) from spatial methods (green). f, Box plots illustrating the distribution of protein-specific imputation accuracy (PCC and SPCC). Statistical significance was determined using a two-sided Wilcoxon signed-rank test: *P<0.05, **P<0.01, ***P<0.001, ****P<0.0001 , ns indicates no significant difference. Center lines indicate medians. g, Robustness analysis of domain identification metrics under varying FOV overlap rates (from 90% down to 10%). h, Robustness analysis regarding the spatial position of the paired window (evaluated across five distinct positions).

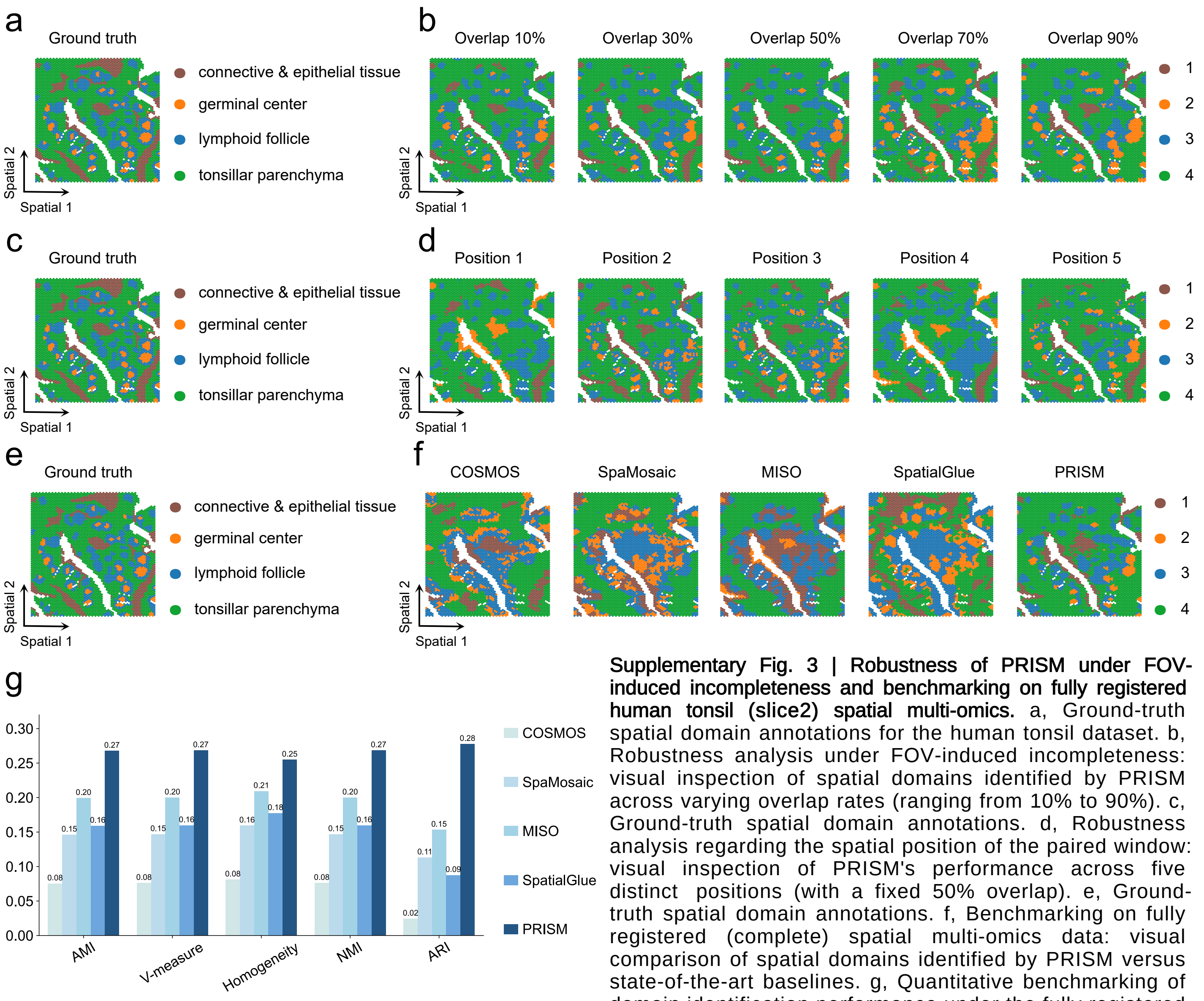


**Supplementary Fig. 3 | Robustness of PRISM under FOV-induced incompleteness and benchmarking on fully registered human tonsil (slice2) spatial multi-omics.** a, Ground-truth spatial domain annotations for the human tonsil dataset. b, Robustness analysis under FOV-induced incompleteness: visual inspection of spatial domains identified by PRISM across varying overlap rates (ranging from 10% to 90%). c, Ground-truth spatial domain annotations. d, Robustness analysis regarding the spatial position of the paired window: visual inspection of PRISM's performance across five distinct positions (with a fixed 50% overlap). e, Ground-truth spatial domain annotations. f, Benchmarking on fully registered (complete) spatial multi-omics data: visual comparison of spatial domains identified by PRISM versus state-of-the-art baselines. g, Quantitative benchmarking of domain identification performance under the fully registered setting using AMI, V-measure, homogeneity, NMI, and ARI.

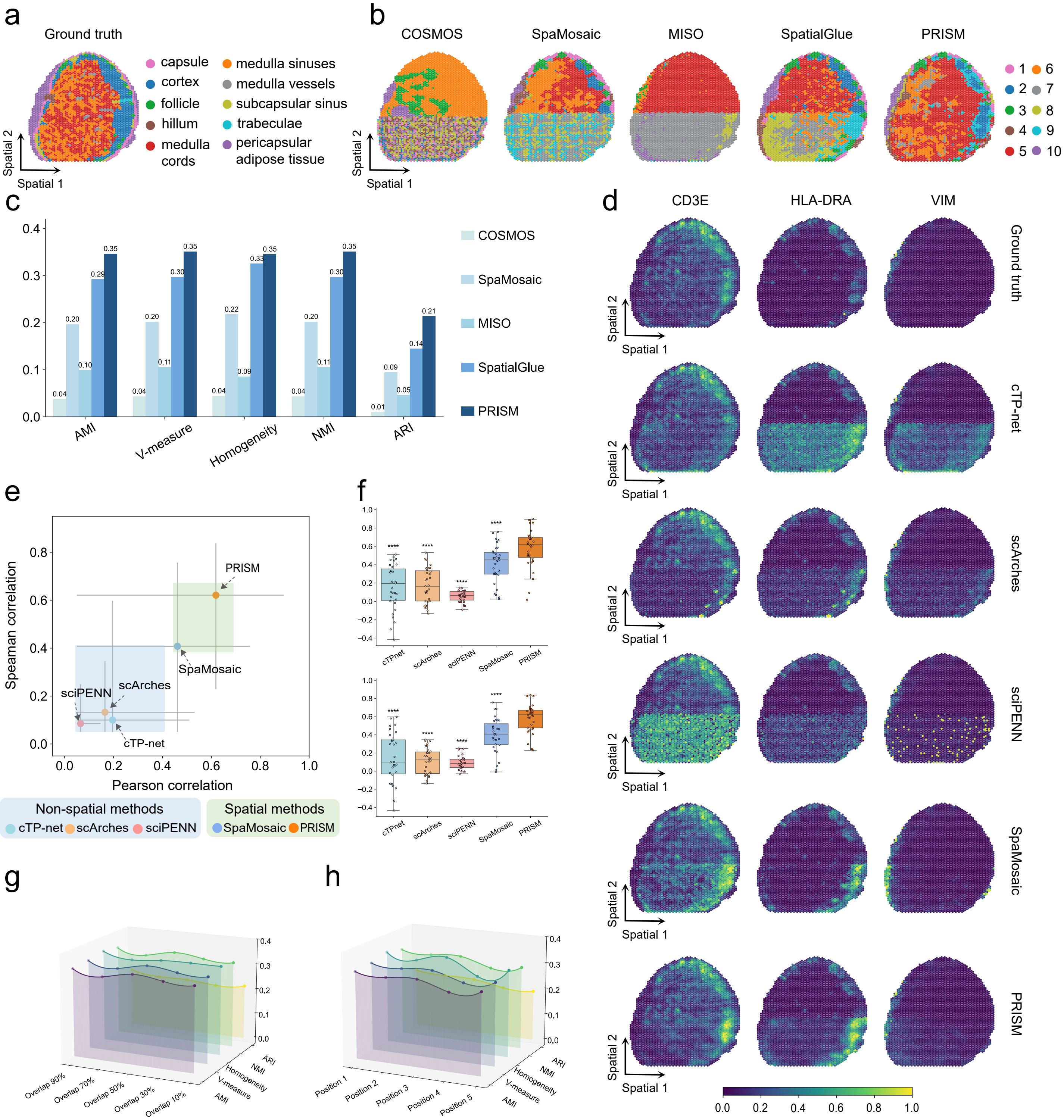


**Supplementary Fig. 4 | Performance of PRISM on human lymph node (slice2) spatial multi-omics under FOV-induced incompleteness.** a,Ground-truth spatial domain annotations for the human lymph node dataset. b, Visual comparison of spatial domains identified by PRISM versus state-of-the-art baselines (COSMOS, SpaMosaic, MISO, and SpatialGlue) under the incomplete spatial multi-omics setting. c,Quantitative benchmarking of domain identification performance using AMI, V-measure, homogeneity, NMI, and ARI. d, Spatial visualization of representative proteins (CD3E, HLA-DRA, and VIM): ground truth measurements compared with imputations from non-spatial translators (cTP-net, scArches, sciPENN) and spatial methods (SpaMosaic, PRISM). e, Scatter plot summarizing overall protein imputation performance (PCC vs. SPCC). Shaded regions distinguish non-spatial methods (blue) from spatial methods (green). f, Box plots illustrating the distribution of protein-specific imputation accuracy (PCC and SPCC). Statistical significance was determined using a two-sided Wilcoxon signed-rank test: *P<0.05 , **P<0.01, ***P<0.001, ****P<0.0001, Center lines indicate medians. g, Robustness analysis of domain identification metrics under varying FOV overlap rates (from 90% down to 10%). h, Robustness analysis regarding the spatial position of the paired window (evaluated across five distinct positions).

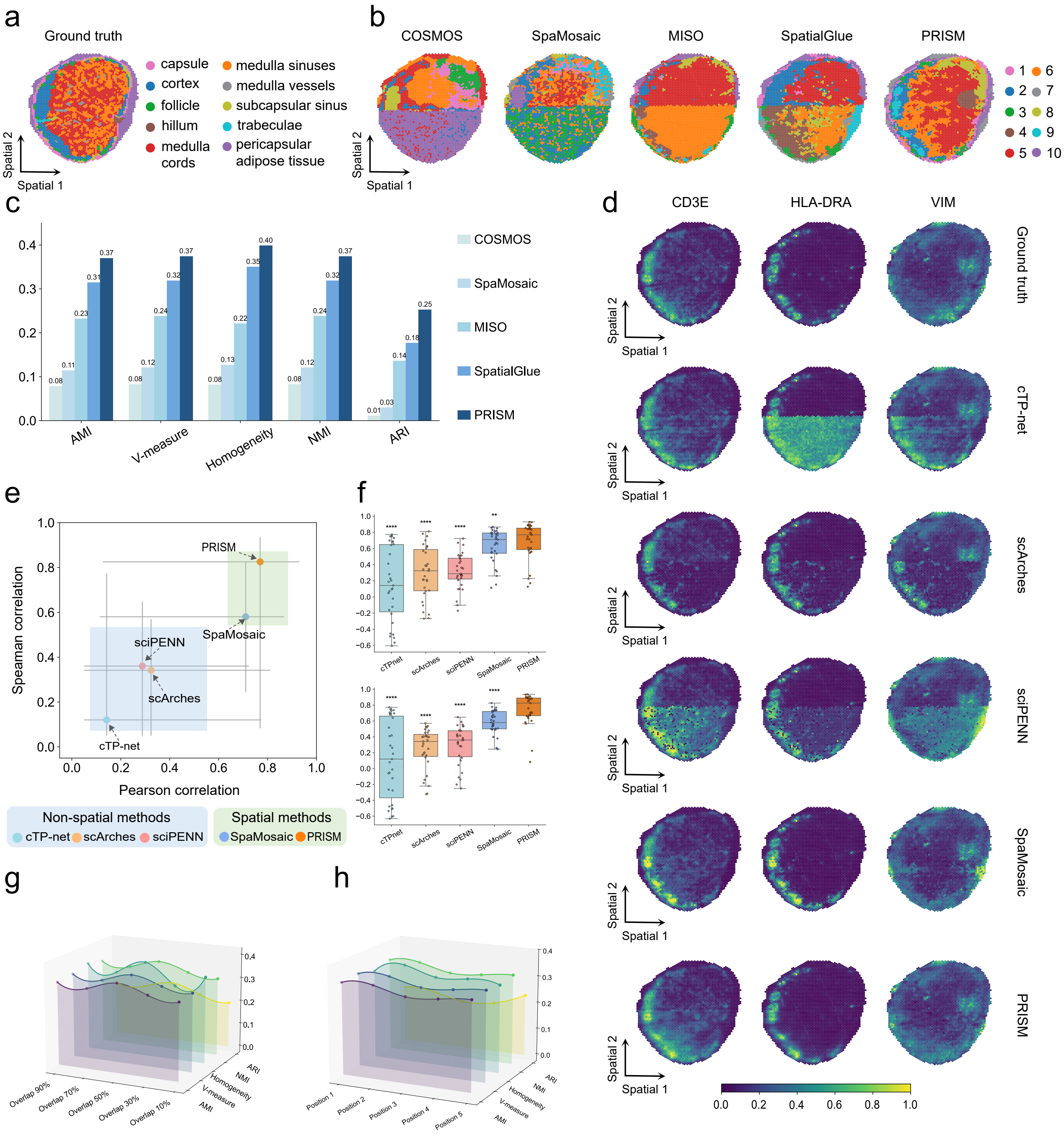

**Supplementary Fig. 5 | Performance of PRISM on human lymph node (slice3) spatial multi-omics under FOV-induced incompleteness.** a,Ground-truth spatial domain annotations for the human lymph node dataset. b, Visual comparison of spatial domains identified by PRISM versus state-of-the-art baselines (COSMOS, SpaMosaic, MISO, and SpatialGlue) under the incomplete spatial multi-omics setting. c,Quantitative benchmarking of domain identification performance using AMI, V-measure, homogeneity, NMI, and ARI. d, Spatial visualization of representative proteins (CD3E, HLA-DRA, and VIM): ground truth measurements compared with imputations from non-spatial translators (cTP-net, scArches, sciPENN) and spatial methods (SpaMosaic, PRISM). e, Scatter plot summarizing overall protein imputation performance (PCC vs. SPCC). Shaded regions distinguish non-spatial methods (blue) from spatial methods (green). f, Box plots illustrating the distribution of protein-specific imputation accuracy (PCC and SPCC). Statistical significance was determined using a two-sided Wilcoxon signed-rank test: *P<0.05, **P<0.01, ***P<0.001, ****P<0.0001, Center lines indicate medians. g, Robustness analysis of domain identification metrics under varying FOV overlap rates (from 90% down to 10%). h, Robustness analysis regarding the spatial position of the paired window (evaluated across five distinct positions).

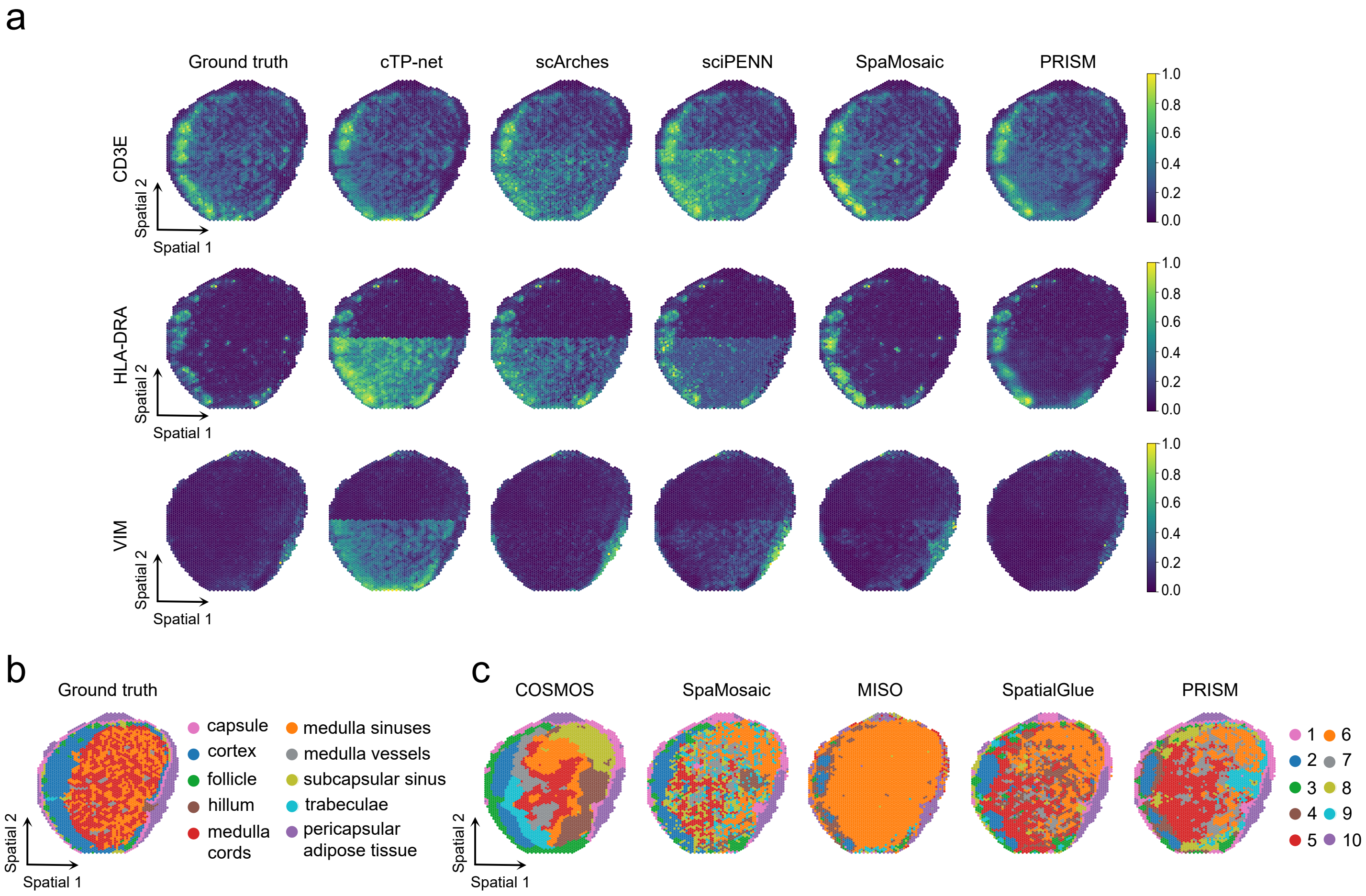


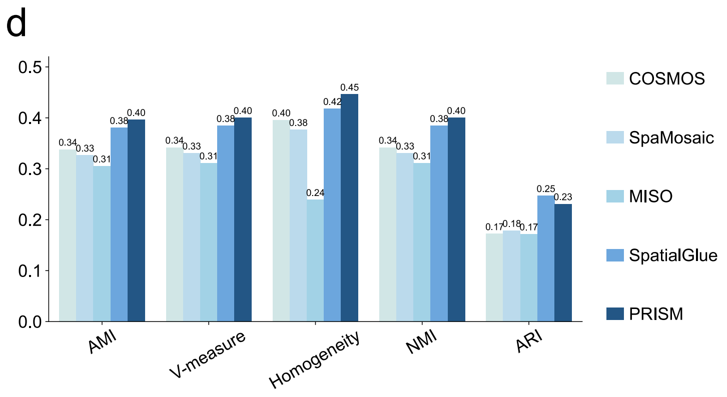


**Supplementary Fig. 6 | Visualization of protein imputation and benchmarking on fully registered human lymph node (slice1) spatial multi-omics.** a, Spatial visualization of representative proteins (CD3E, HLA-DRA, and VIM): comparison of imputations from non-spatial translators and spatial methods. b, Ground-truth spatial domain annotations. c, Benchmarking on fully registered (complete) spatial multi-omics data: visual comparison of spatial domains identified by PRISM versus state-of-the-art baselines. d, Quantitative benchmarking of domain identification performance under the fully registered setting using AMI, V-measure, homogeneity, NMI, and ARI.

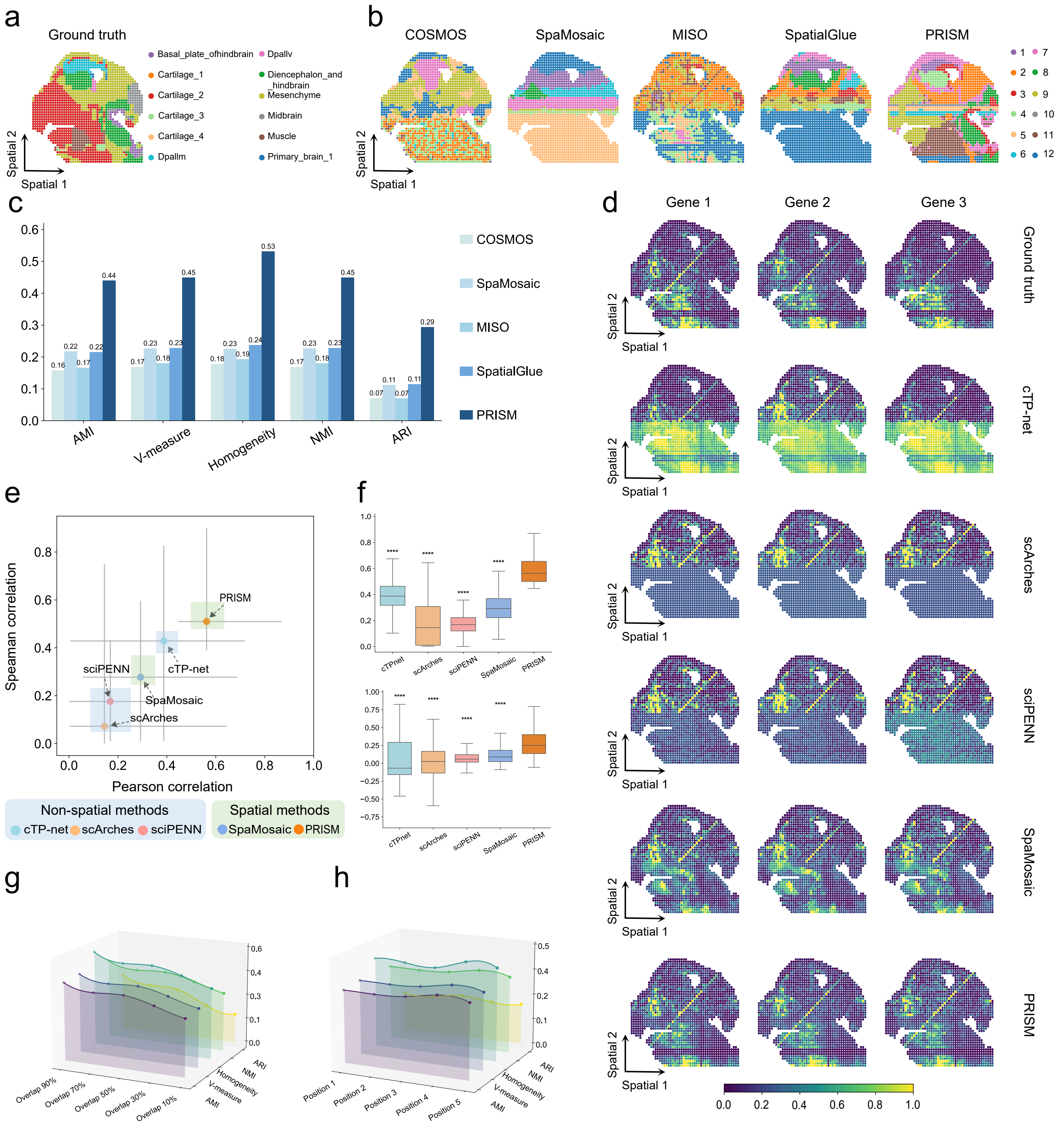


**Supplementary Fig. 7 | Performance of PRISM on mouse embryonic brain (slice1) spatial multi-omics under FOV-induced incompleteness.** a, Ground-truth spatial domain annotations for the mouse embryonic brain dataset (including basal plate of hindbrain, cartilage, Dpallv, diencephalon and hindbrain mesenchyme, midbrain, muscle, and primary brain regions). b, Visual comparison of spatial domains identified by PRISM versus state-of-the-art baselines under the incomplete spatial multi-omics setting. c, Quantitative benchmarking of domain identification performance using AMI, V-measure, homogeneity, NMI, and ARI. d, Spatial visualization of representative genes: ground truth measurements compared with imputations from non-spatial translators (cTP-net, scArches, sciPENN) and spatial methods (SpaMosaic, PRISM). e, Scatter plot summarizing overall gene imputation performance (PCC vs. SPCC). Shaded regions distinguish non-spatial methods (blue) from spatial methods (green). f, Box plots illustrating the distribution of gene-specific imputation accuracy (PCC and SPCC). Statistical significance was determined using a two-sided Wilcoxon signed-rank test: *P<0.05, **P<0.01, ***P<0.001, ****P<0.0001. Center lines indicate medians. g, Robustness analysis of domain identification metrics under varying FOV overlap rates (from 90% down to 10%). h, Robustness analysis regarding the spatial position of the paired window (evaluated across five distinct positions).

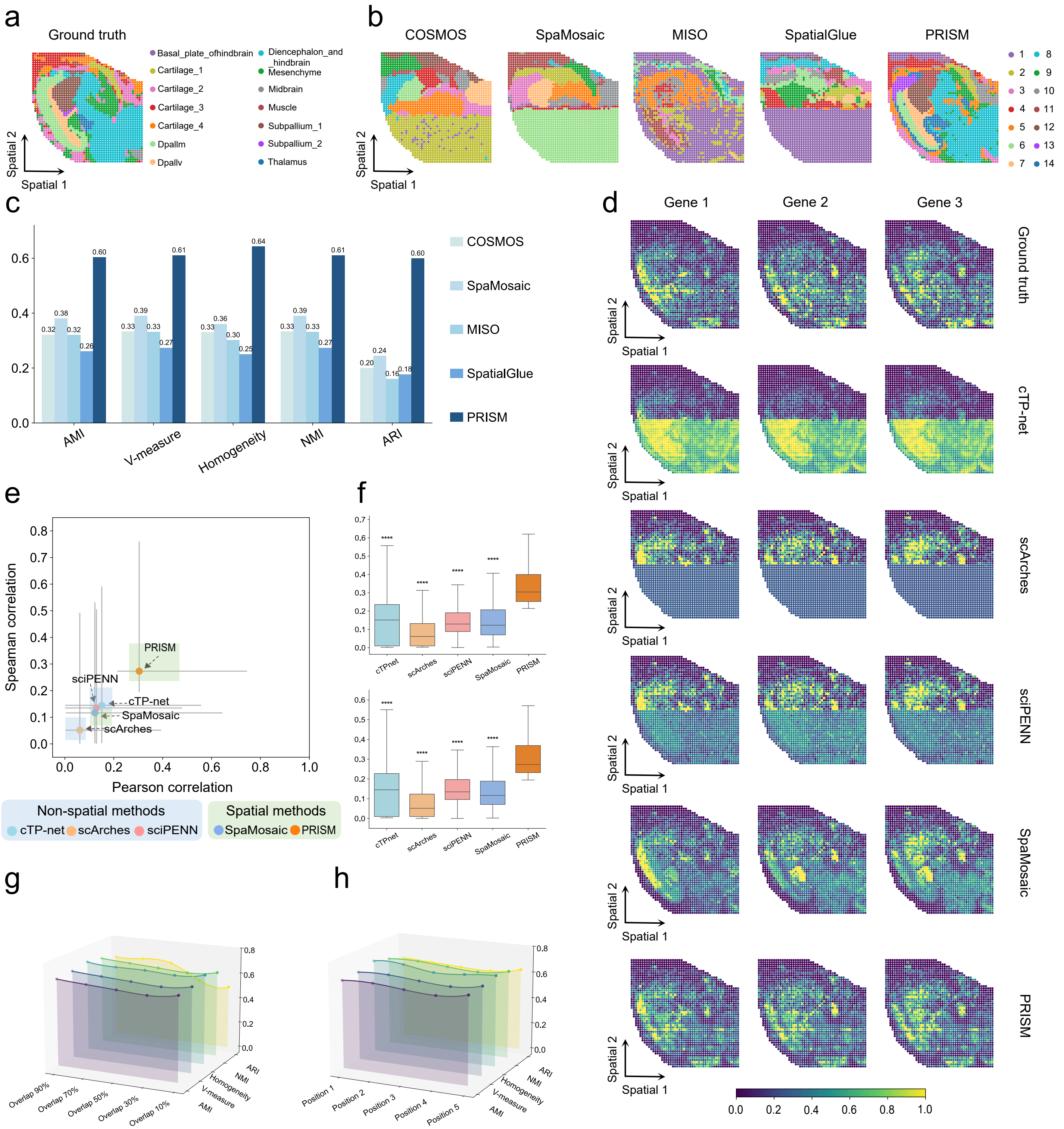


**Supplementary Fig. 8 | Performance of PRISM on mouse embryonic brain (slice3) spatial multi-omics under FOV-induced incompleteness.** a, Ground-truth spatial domain annotations for the mouse embryonic brain dataset. b, Visual comparison of spatial domains identified by PRISM versus state-of-the-art baselines under the incomplete spatial multi-omics setting. c, Quantitative benchmarking of domain identification performance using AMI, V-measure, homogeneity, NMI, and ARI. d, Spatial visualization of representative genes: ground truth measurements compared with imputations from non-spatial translators (cTP-net, scArches, sciPENN) and spatial methods (SpaMosaic, PRISM). e, Scatter plot summarizing overall gene imputation performance (PCC vs. SPCC). Shaded regions distinguish non-spatial methods (blue) from spatial methods (green). f, Box plots illustrating the distribution ofgene-specific imputation accuracy (PCC and SPCC). Statistical significance was determined using a two-sided Wilcoxon signed-rank test: *P<0.05, **P<0.01, ***P<0.001, ****P<0.0001. Center lines indicate medians. g, Robustness analysis of domain identification metrics under varying FOV overlap rates (from 90% down to 10%). h, Robustness analysis regarding the spatial position of the paired window (evaluated across five distinct positions).

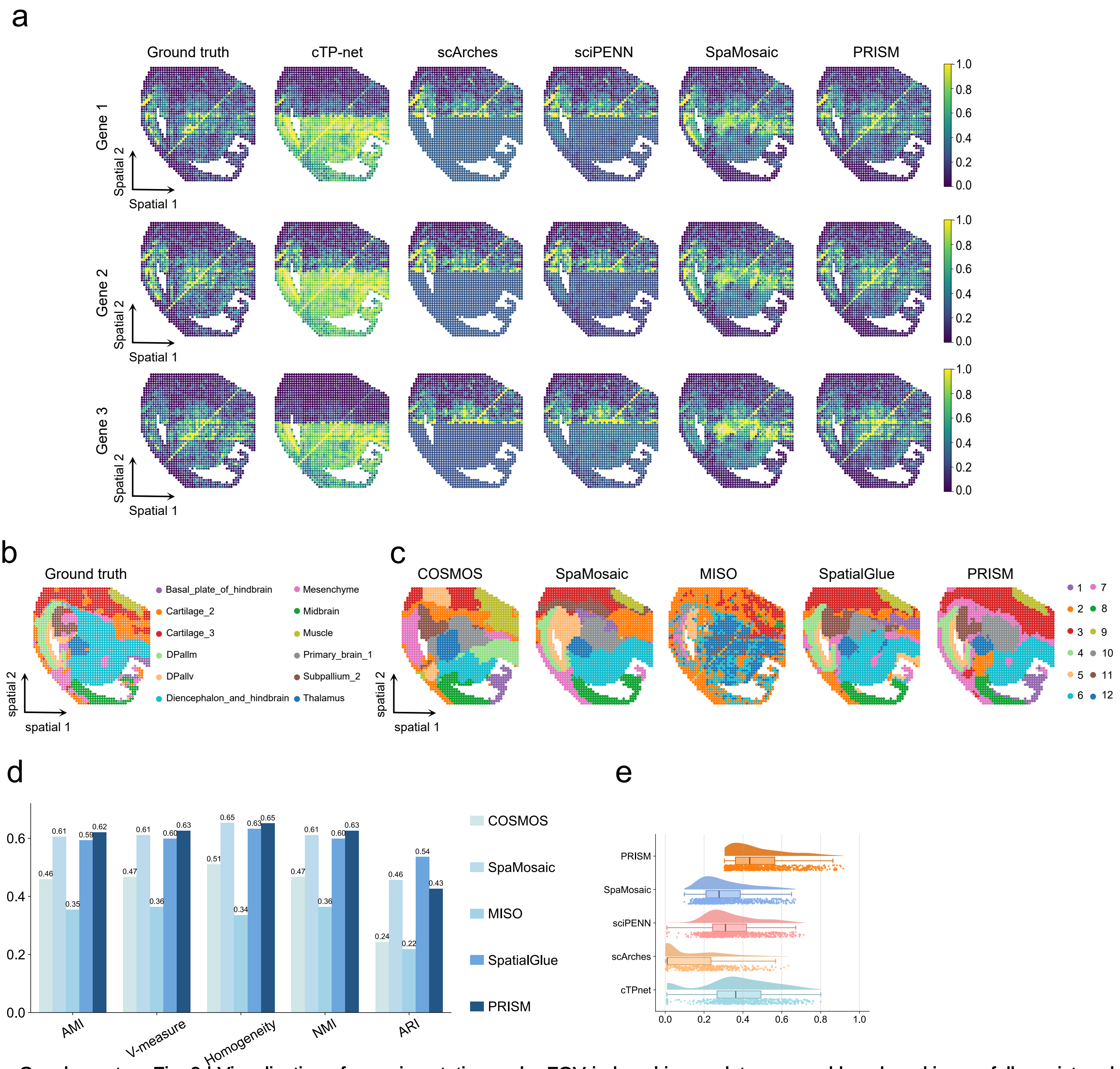


**Supplementary Fig. 9 | Visualization of gene imputation under FOV-induced incompleteness and benchmarking on fully registered mouse embryonic brain (`slice2`) spatial multi-omics.** a, Spatial visualization of representative genes under FOV-induced incompleteness: comparison of ground truth measurements with imputations from non-spatial translators (cTP-net, scArches, sciPENN) and spatial methods (SpaMosaic, PRISM). b, Ground-truth spatial domain annotations for the mouse embryonic brain dataset. c, Benchmarking on fully registered (complete) spatial multi-omics data: visual comparison of spatial domains identified by PRISM versus state-of-the-art baselines (COSMOS, SpaMosaic, MISO, and SpatialGlue). d, Quantitative benchmarking of domain identification performance under the fully registered setting using AMI, V-measure, homogeneity, NMI, and ARI. e, Distribution of imputation accuracy (SPCC) for the top 800 highly variable genes (HVGs) under the incomplete setting. Center lines indicate medians, points represent individual genes.

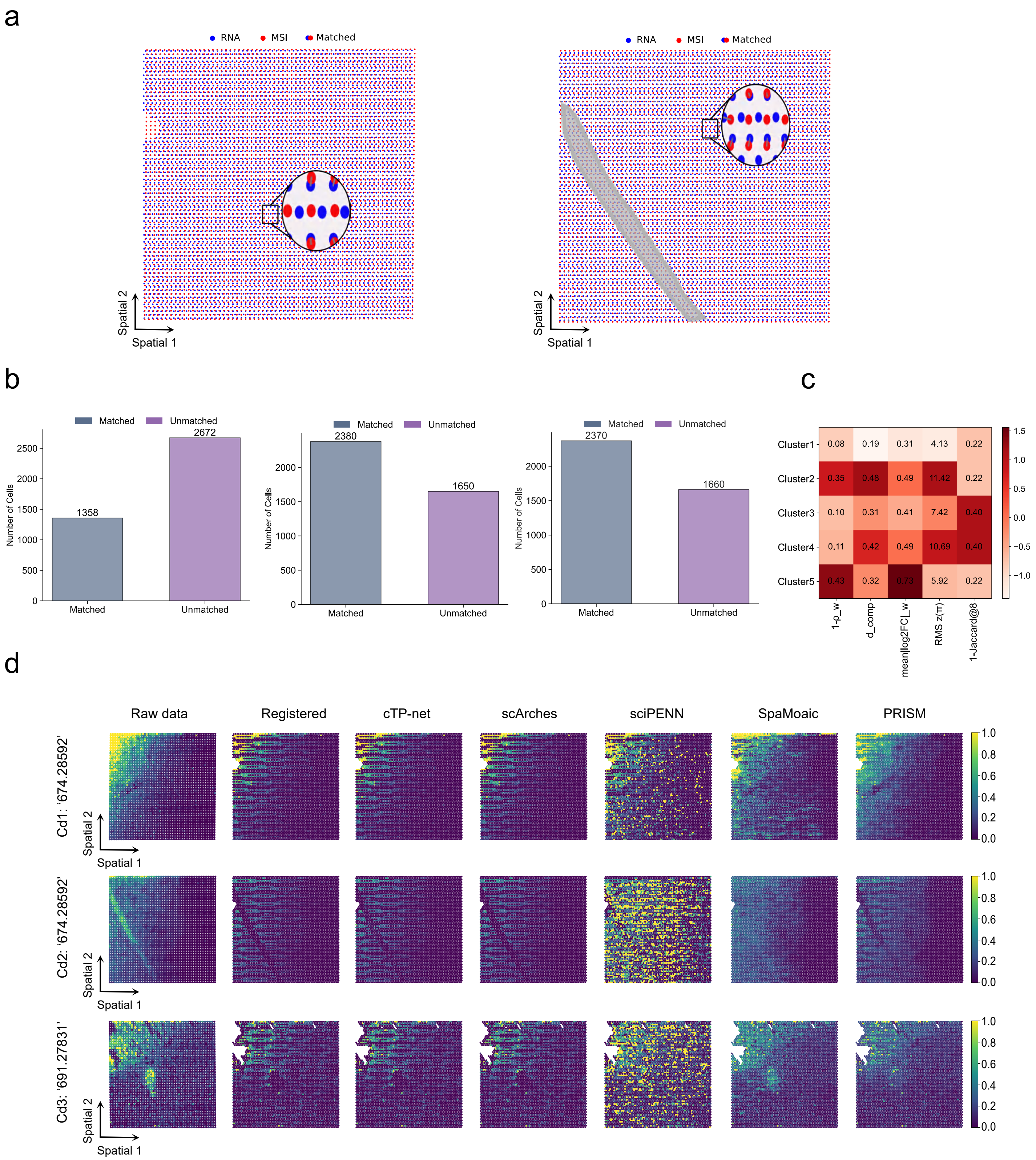


**Supplementary Fig. 10 | Characterization of resolution-induced incompleteness and performance evaluation in human PD striatum.** a, Visualization of cross-modality alignment for slices Cd1 and Cd2, highlighting the spatial distribution of RNA spots (blue), MSI pixels (red), and matched pairs. The zoom-in views illustrate the spatial disparity and resolution-induced incompleteness between the two modalities. b, Bar plots quantifying the number of matched versus unmatched spatial spots across three slices (Cd1 – Cd3), explicitly demonstrating the extent of data sparsity resulting from the registration process. c, Quantitative assessment of differential expression quality across identified spatial domains (Clusters 1-5), evaluated using multiple validity metrics (including p-value significance, fold change, and specificity). d, Spatial visualization of representative metabolites (m/z 674.28592, m/z 691.27831) across three slices. Columns compare: (1) Raw data (unfiltered, showing original signals before registration masking), (2) Registered input (showing resolution-induced gaps where signals were treated as missing), and (3) Imputations from non-spatial translators (cTP-net, scArches, sciPENN) and spatial methods (SpaMosaic, PRISM). PRISM effectively restores the signal continuity and intensity patterns observed in the raw data, whereas baselines exhibit artifacts or signal loss.

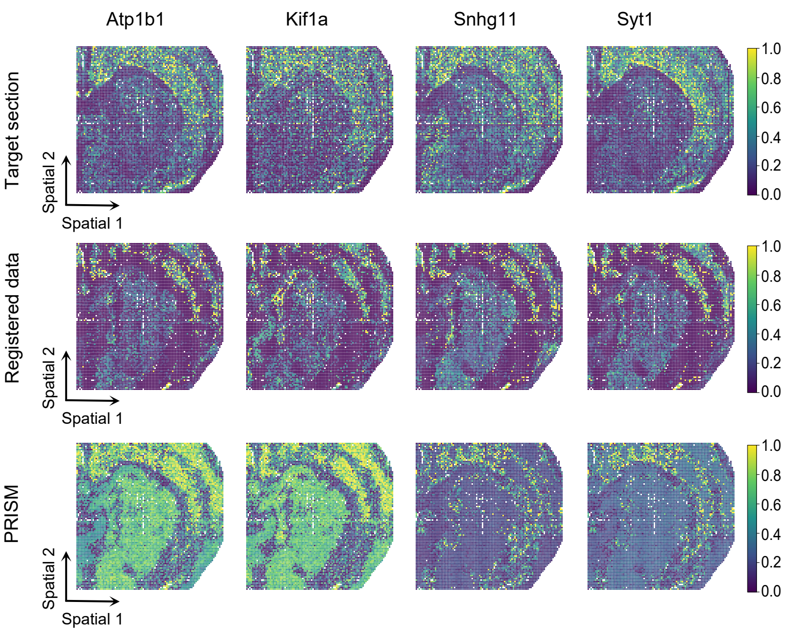


**Supplementary Fig. 11 | Visualization of cross-slice gene imputation in adjacent-slice P22 mouse brain.** Spatial visualization of four representative neuronal marker genes (Atp1b1, Kif1a, Snhg11, and Syt1) across three conditions. Top row (Target section): The original, ground-truth gene expression observed in the target transcriptomic slice. Middle row (Registered data): The input data after adjacent-slice registration (via SLAT) and quality filtering. This panel highlights the incomplete spatial multi-omics nature of the input, characterized by significant sparsity and signal discontinuity due to the exclusion of low-confidence matches. Bottom row (PRISM): The imputed gene expression profiles generated by PRISM. PRISM effectively reconstructs the continuous spatial patterns and laminar gradients (e.g., in Syt1) that were fragmented in the registered input, closely mirroring the anatomical structures of the target section.
